# Linking intrinsic scales of ecological processes to characteristic scales of biodiversity and functioning patterns

**DOI:** 10.1101/2021.10.11.463913

**Authors:** Yuval R. Zelnik, Matthieu Barbier, David W. Shanafelt, Michel Loreau, Rachel M. Germain

## Abstract

Ecology is a science of scale, which guides our description of both ecological processes and patterns, but we lack a systematic understanding of how process scale and pattern scale are connected. Recent calls for a synthesis between population ecology, community ecology, and ecosystem ecology motivate the integration of phenomena at multiple levels of organization. Furthermore, many studies leave out the scaling of a critical process: species interactions, which may be non-local through movement or foraging and must be distinguished from dispersal scales. Here, we use simulations to explore the consequences of three different process scales (species interactions, dispersal, and the environment) on emergent patterns of biodiversity, ecosystem functioning, and their relationship, in a spatially-explicit landscape and stable equilibrium setting. A major result of our study is that the spatial scales of dispersal and species interactions have opposite effects: a larger dispersal scale homogenizes spatial biomass patterns, while a larger interaction scale amplifies their heterogeneity. Interestingly, the specific scale at which dispersal and interaction scales begin to influence landscape patterns depends on the scale of environmental heterogeneity – in other words, the scale of one process allows important scales to emerge in other processes. This interplay between process scales, i.e., a situation where no single process dominates, can only occur when the environment is heterogeneous and the scale of dispersal small. Finally, contrary to our expectations, we observe that the spatial scale of ecological processes is more clearly reflected in landscape patterns (i.e., distribution of local outcomes) than in global patterns such as Species-Area Relationships or large-scale biodiversity-functioning relationships. Overall we conclude that long-range interactions often act differently and even in opposite ways to dispersal, and that the landscape patterns that emerge from the interplay of long-ranged interactions, dispersal and environmental heterogeneity are not well captured by often-used metrics like the Species-Area Relationship.

## Introduction

Scale is fundamental to ecology, from the spatial and temporal scales at which we observe and manage ecosystems [1, 2, 3] to the intrinsic scales at which processes occur within and across ecosystems [4]. Much of current research efforts describe ecological patterns across scales, such as Species-Area or Biodiversity-Ecosystem Functioning relationships [5, 3]. However, the scaling of ecological patterns is largely phenomenological – we can describe how patterns scale but not why [6, 5]. Although links between scales of patterns and processes have been explored in recent years [7, 8, 9], as we will discuss, a systematic and unified treatment of scale in ecology is incomplete. A critical question remains: how is the scaling of ecological patterns, such as patterns of biodiversity and ecosystem functioning, generated by scales of specific processes, and why?

In answering this question, a crucial process is often overlooked: the spatial scale of species interactions. While dispersal and environmental variation are often understood to operate at various spatial scales, existing research generally assumes that species only interact locally [10, 11, 12] (although exceptions exist, e.g., studies using multi-layer networks to link interaction networks at local scales to their realization at the global scale [13, 14]). Yet many species move, forage, or otherwise interact with each other at a range of spatial scales [15, 16], even in the absence of dispersal. A simple distinction is that dispersing species establish new “home” ranges when they move across the environment, while mobile species always return to their “home” range. Many move daily across multiple habitat types, such as seabirds connecting marine and terrestrial ecosystems [15], or predatory insects moving between different habitats in the landscape [16]. Non-local competition can therefore arise from foraging across multiple localities. Additionally, species interact indirectly across long distances via intermediary species, (e.g., plants interacting indirectly via pollinators or herbivores), and many such intermediary interactions are not explicitly studied, thus being best represented by long range interactions. As a result, scales of species interactions, such as competition, likely have consequences for population persistence, affecting the spatial distribution of biodiversity and ecosystem functioning in ways that are distinct from other process scales [17, 18].

How do the spatial scales of dispersal, environmental heterogeneity, and species interactions interactively influence ecological patterns? Answering this question is unlikely to be achieved via observational studies, as different combinations of ecological processes may generate identical patterns, but computational models can explore patterns that emerge as processes interact across scales. Indeed, the scale of dispersal relative to the environment has been studied most extensively, in particular within a metacommunity context [19, 7, 20]. These studies generally find that high rates of dispersal blur differences between local communities, leading to losses of biodiversity and ecosystem functioning. Although there are reasons to expect increased scales of dispersal and species interactions to have similar consequences, as both processes are influenced by many of the same variables (e.g., animal mobility) and serve to spread out the effects of species interactions, there are also reasons to expect the opposite [21]. A key difference is that large dispersal scales can allow populations to permeate through whole landscapes over a few generations, whereas individuals with large interaction scales are still bound to specific localities. As a result, increasing scales of interactions may amplify spatial heterogeneity in an ecological system [22], counter to the blurring effect of larger dispersal scales.

In addition to scales of species interactions, we will address an additional major gap which prevents a complete knowledge of scaling in ecology: consideration of a wider range of ecological patterns within a single study than has been examined previously. Two well-recognized ecological patterns are Species-Area (SAR) and Biodiversity-Ecosystem Functioning (BEF) relationships. The Species-Area relationship is the earliest and most widely-examined ecological pattern to explicitly consider scale [5, 23]. Although SARs have been described as one of “ecology’s few universal regularities” [24], accumulating evidence reveals considerable variation within and among biological systems [25, 5, 26]. Likewise, BEF theory has revealed consistent patterns, typically a saturating relationship between community diversity and biomass production [27], but most work has focused on BEFs at local scales, with only recent work highlighting the importance of scale [3]. Previous studies have examined how one pattern or the other are affected by process scales [28, 26, 29], but no study has examined how SAR and BEF relationships change in tandem and if effects that are masked through one pattern are apparent in the other. As a consequence, it is unclear how both SAR and BEF relationships are affected by the interplay of processes acting at different scales, making it difficult to assess how process scales affect the overall behavior of ecosystems as different measures highlight different aspects of ecosystems. Resolving these issues will be useful for both basic and applied biodiversity problems, for instance allowing us to scale up to landscape scales our predictions of biodiversity loss and its effect of ecosystem productivity, that are often based on local scales [30].

Here, we use a modified Lotka-Volterra metacommunity model to explore the consequences of the scaling of ecological processes for biodiversity, ecosystem functioning, and their relationship across spatial scales. Our simulations consist of species interacting in a spatially-explicit landscape, with “patches” emerging from the environmental structure of the landscape. Although metacommunities tend to be modelled as systems of discrete patches embedded within an inhospitable matrix, Chase and Leibold [31] describe this approach as useful (easing computation and interpretation) but limited – they foreshadow a “coming” in ecology in favour of models that allow “patches” to emerge from the structure of the environment, which our model achieves. We first study the heterogeneity of local outcomes across the landscape: patterns of patch biodiversity, patch functioning, and relationships between them (local BEF). We can then scale up to the whole landscape scale and every scale in between. By varying the spatial scales over which metacommunity processes (abiotic environment, competitive interactions, and dispersal) play out, we test the hypothesis that ecological patterns depend on how processes interact across scales, including scales of species interactions, and lead to different patterns from those generated by commonly-assumed hierarchical process scales (i.e., scales of interactions *<* environment *<* dispersal; Fig. 1).

**Figure 1:**
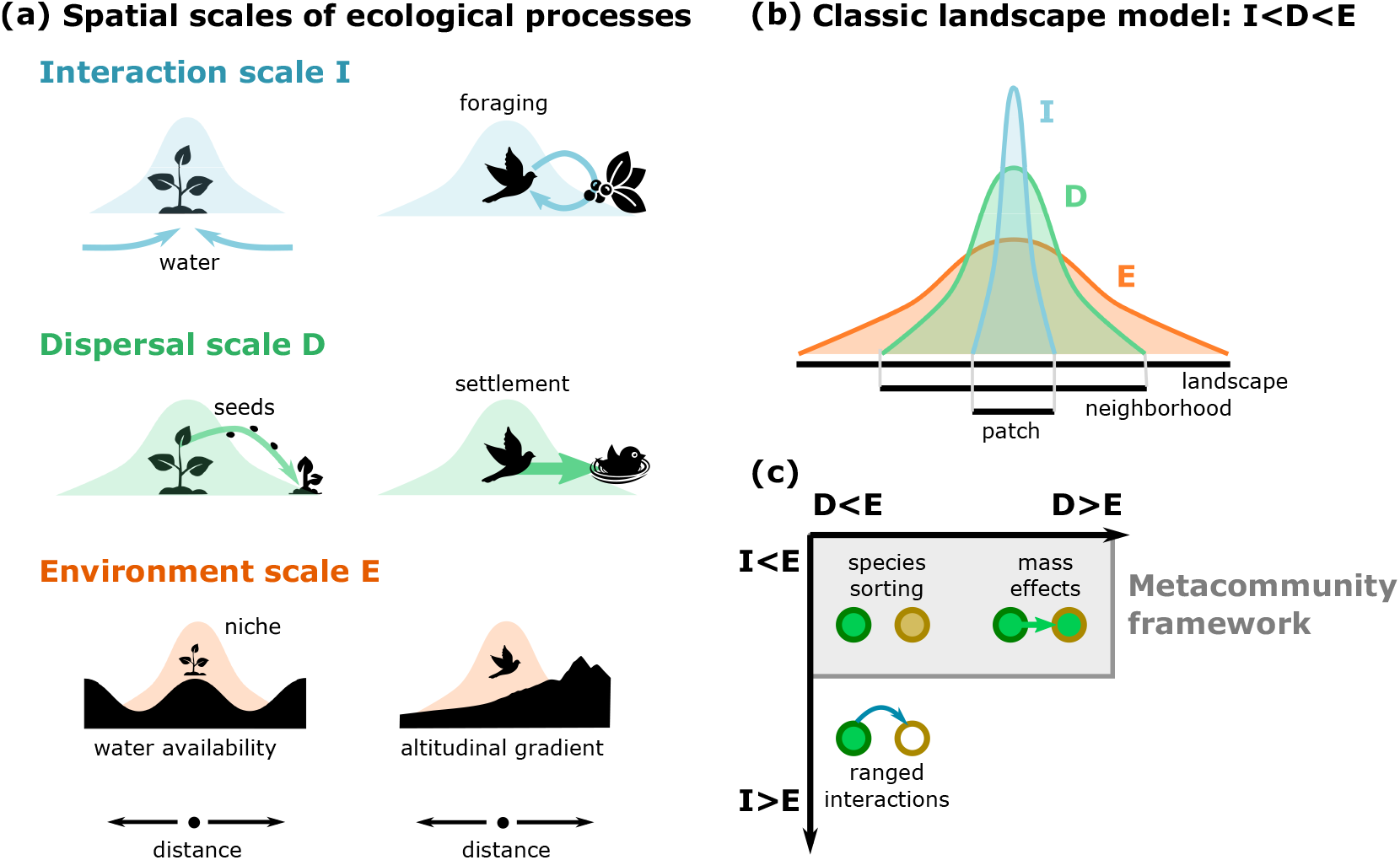
Conceptual diagram of spatial scales of ecological processes. **(a)** Illustration of the spatial scale of species interactions I, dispersal D and environmental heterogeneity E relative to the total size of the landscape (i.e., width of curves). **(b)** In the classic scenario, interactions take place within a patch, while dispersal is thought to act within a neighborhood and environmental factors vary broadly over the landscape. **(c)** Comparison of ecological scenarios along scales of I, D and E. Yellow and green represent two different species, with circle and its rim representing the resident species and the favoured species, respectively. Metacommunity theory has explored different scenarios for the relative scales of dispersal and environment (i.e., the ratio D/E), notably distinguishing “species sorting” (local environmental factors determine species distribution) and “mass effects” (population fluxes homogenize the landscape). Our work highlights the relative importance of species interactions scale (e.g., expressed through the ratio I/E, which was previously considered only in particular ecological settings (e.g., vegetation patterns or territoriality). Ranged interactions may for instance induce exclusion of weaker competitors in a neighboring patch, even without a population flux of a stronger competitor into that patch.

Species-Area relationships depend on spatial turnover in species composition, and compositional turnover is driven by ecological processes [32]. Thus, we would expect that ecological processes should strengthen SARs in scenarios where they increase compositional turnover. We predict that the strongest slopes of the SAR will occur when scales of dispersal *<* environment *<* species interactions, because (i) interactions are not constrained to abiotically suitable patches, and (ii) weaker dispersal prevents the homogenization of species composition across the landscape. Additionally, we predict that the consequences for BEF relationships will differ between local and regional scales. On local scales, we expect BEFs to weaken as interaction scales increase relative to the others, given that species that are locally absent but present in nearby areas can affect local functioning. On regional scales, we expect BEFs to strengthen as interaction scales increase, since regional competition would keep only the most suitable species at a given location. Hence, more species would mean that multiple species are productive within a given region.

## Methods

### Model

We use a modified Lotka-Volterra metacommunity model to explore the consequences of the spatial scaling of three ecological processes – abiotic environment, species interactions, and dispersal – for biodiversity and ecosystem functioning. Our specific assumptions and parameters are motivated by two important choices. First, we focus on a classic setting of ecological assembly, i.e., the patterns that arise when many species, originating from a regional pool, come together and reach an equilibrium state, with some species going locally or regionally extinct. Furthermore, we take species interactions in the pool to be disordered, that is, heterogeneous but without a particular functional group or trophic level structure [33]. We do not exclude that different patterns could emerge for more ordered interactions (e.g., a realistic food web) or for parameter values that lead to more complex dynamical regimes (e.g., population cycles or chaos, driven by stronger species interactions or environmental perturbations). We note that our communities, in the chosen parameter regime of moderate competition, contain many species in a stable equilibrium (i.e., due to the assembly process). Our methodology thus differs from the extensive literature that has considered models with random interactions in order to study stability-complexity relationships [34], including more recent works in a spatial context [35, 36], as we rather focus on the abundance and diversity patterns arising from community assembly.

Second, we consider the possibility of species interacting over large spatial scales. Conventional metacommunity models describe discrete local communities of habitat patches connected by dispersal, within which species interact [37]. In doing so, they implicitly assume that the spatial range of species interaction is smaller than the scale of dispersal and contained within a patch, for all species and types of interactions [17]. To relax these assumptions, we construct a metacommunity model where populations of species can disperse and interact at different spatial scales, without specifying a mechanism underlying these ecological processes. Species interactions that manifest beyond local scales are abstracted from mechanisms such as individual foraging, vector species (e.g., pathogens) [38], and spatial resource fluxes [39, 17].

The model details the dynamics of *S* different species distributed across a spatially-explicit lattice landscape of 320×320 cells. The dynamical equation for the biomass *N*_*i*_ of species *i* at position 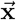 in the landscape at time *t* is given by a generalized Lotka-Volterra equation of the form

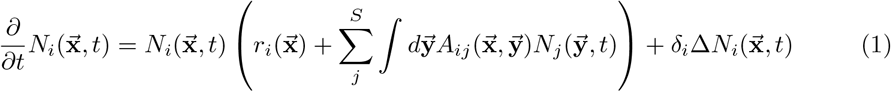

where 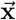 and 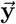 represent vectors of spatial (*x, y*) coordinates in the landscape. Equation (1) models the effects of three ecological processes on the biomass of species *i*: its intrinsic growth rate 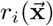, which is influenced by abiotic environmental conditions at location 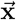, dispersal to and from location 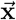, which is controlled by the diffusion coefficient *δ*_*i*_, and interactions with all other species *j*, including when they are located elsewhere in the landscape, 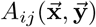. Although at face value cells in our model resemble patches in traditional metacommunity models, given that discrete populations are necessary to simulate Lotka-Volterra dynamics, here it is best to interpret cells as neighborhoods on a landscape. Each neighborhood may take on a unique environmental value and hold unique densities of individuals of different species. Viewed in this way, landscape dynamics can be simulated more continuously, with the numerical limitation of needing to discretize dynamics at their finest resolution. While “patches” can emerge in autocorrelated environments (i.e., a spatial clustering of cells that are suitable to a given species), our model is also generalizable to landscapes with a diversity of environmental structures.

### Environment

Abiotic conditions in each location are encoded by an environmental variable 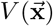. This variable is continuous and varies smoothly over space, with parameters allowing one to tune the typical spatial scale of this variation [40]. For more details on the construction of the environment, see the Appendix section A2.

Each species has a Gaussian fundamental niche that determines its abiotic fitness in each location, with an optimal environmental value *H*_*i*_ and abiotic niche width *ω*_*i*_

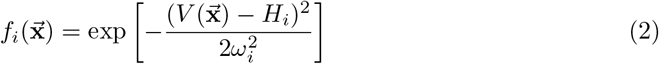

Each fitness value is bound between 0 and 1 and reaches its maximum at an optimal environmental condition (i.e., when 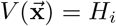). We take the growth rate as 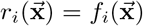. In other words, 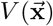 sets the actual structure of environmental conditions across the landscape, whereas 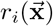 is how species experience the environment and its structure.

### Interactions

We choose to limit ourselves to competitive interactions, defined by the matrix *C*_*ij*_, which represents the per-capita competitive effect of species *j* on species *i*. The diagonal of the matrix (the impact of a species on itself) is set to 1, whereas all other interactions are taken independently from a random uniform distribution between 0 and 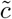. We choose 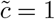 to allow for moderate interactions between different species (inter-specific competition is always weaker than intra-specific), suggesting that pairwise coexistence is often possible for species with different growth rates *r*_*i*_, but the total impact of many competitors is still strong enough to allow for extinctions. Previous work has shown that, in disordered communities, the outcomes of ecological assembly are robust to many details such as the nature of interactions (e.g., mutualism, predation), and depend only on a few statistical properties such as the mean and variance of interaction effects [33].

Furthermore, interactions are assumed to occur over a characteristic spatial scale encoded by a spatial kernel *K*. This scale may represent the distance an animal forages from its nest (without establishing a new nest), the scale at which trees gather resources with their roots, or the effective distance an immobile species interacts with its neighbors via an intermediary species (where the intermediary is not explicitly modeled). We use a Gaussian kernel whose standard deviation defines the interaction range such that

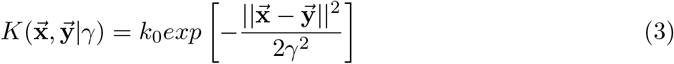

where 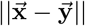 indicates the norm of (distance between) the vectors 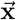 and 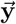, and *γ* is the spatial range (scale) of the interactions. We note that while this modeling strategy is not physical as it implies that interactions occur instantaneously across distances, this is not expected to bias our results since we are focusing on the equilibrium state of the system, where hypothetical lag effects should be minimal.

We normalize the interactions by *k*_0_ such that the overall effect of the kernel is always the same (i.e., the integral over K always equals 1). This normalization means that for large-scale interactions, local competition becomes weaker. However, some amount of (especially intra-specific) competition must remain locally strong to prevent species densities from growing exponentially and exploding. Therefore, we define interactions as partially local and partially regional, with *β* governing the fraction of interactions that are regional:

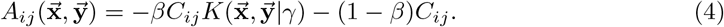

We choose *β* to ensure that the effect of interactions changes with their spatial scale (see scales subsection below), but local competition is never negligible (see more details in the Appendix, Fig. S12).

### Dispersal

Finally, dispersal is modeled by the diffusion (Laplace) operator,

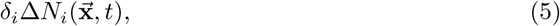

where *δ*_*i*_ is the diffusion or dispersal coefficient of the species. For simplicity, we set the dispersal coefficient to be the same for all species.

Contrary to interactions, we do not use an explicit spatial kernel here, because intensity and spatial scale are unavoidably entangled in the case of dispersal (see Appendix section A1). The coefficient *δ*_*i*_ sets the spatial scale over which dispersal impacts ecological dynamics. Note that two aspects of our modeling choices mean that our choice of dispersal by diffusion is not qualitatively different from applying a large dispersal kernel: our focus on the equilibrium state, and having initial conditions where all species are introduced to every point in the landscape. The former aspect of equilibrium means that any potential non-equilibrium dynamics driven by species moving quickly across space due to a large dispersal kernels are not applicable. The latter aspect means that there is no limit to dispersal, i.e., a short or long-ranged dispersal kernel does not affect which parts of the landscape can be reached by a species.

### Scales

In this study we are concerned with spatial scales of three ecological processes:

1. *E*: environmental heterogeneity
2. *D*: dispersal
3. *I*: species interactions

To properly compare the interplay of different process scales, we must first compute their values for a given set of model parameters (Table 1). The scale of the environment combines two features often used in the literature to generate realistic, spatially-autocorrelated landscapes [41]: spectral color *ρ*, which indicates the relative importance of long-range and short-range variations in the environment, and spectral cutoff *k*_*c*_, which indicates the finest grain of variation (Appendix section A2). The effective environmental scale *E* is controlled by these two parameters.

**Table 1:** Parameters, default values and ranges.

| Parameter | Interpretation | Baseline value [Range] |
| --- | --- | --- |
| **General** |  |  |
| $S$ | species number | 20 |
| $L$ | landscape size (cells) ($area = L^2$) | 320 |
| $\delta_i$ | dispersal coefficient | [0.01, 100] |
| **Environment** |  |  |
| $H_i$ | optimal environment value | $\sim \text{uniform}(20, 80)$ |
| $\omega_i$ | abiotic niche width | $\sim \text{normal}(10, 2)$ |
| $\rho$ | spectral color | 0.95 |
| $k_c$ | spectral cutoff | 0.04 |
| $K(\vec{x})$ | local abiotic conditions | [0, 100] |
| $k_0$ | normalization constant | - |
| **Interactions** |  |  |
| $\tilde{c}$ | max interaction strength | 1.0 |
| $\beta$ | fraction of regional interactions | 0.9 |
| $\gamma$ | spatial scale of interactions | [1, 100] |
| $C_{ij}$ | interaction matrix | $\sim \text{uniform}(0, \tilde{c})$ |

In the main text, we focus on a single value for the environment scale *E* = 32, and vary the other two scales on a logarithmic scale, with values of 1, 3.2, 10, 32 and 100, where the system itself has the scale (length) of 320 cells. Our distribution of *I* and *D* are equally spaced along a log scale and allow us to have a clear separation between the scales of each ecological process, while also being substantially smaller than the system size (320 cells) and larger than the smallest scale in the system (1 cell). Details on the construction of the environment are given in the Appendix section A2. We choose a value of *E* = 32 specifically as it is the most straightforward to demonstrate our results (see Appendix section A3 for other values). The scale of interactions is set by, and coincides with, the width of the Gaussian kernel *γ*, such that *I* = *γ*. The scale of dispersal is <u>m</u>ainly determined by the diffusion coefficient *δ*_*i*_, and it is expected to scale as 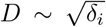 (see, e.g., [42]). The normalization constant is, however, not trivial, and as w<u>e</u> show in the Appendix section A1, it is approximately 10. We therefore use: 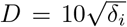. Fixing the environmental scale and varying the scale of interactions and dispersal allows us to isolate the effects of interaction and dispersal scale without confounding the effects of different landscape structures or differences between species.

### Parameterization and simulations

To initialize our simulations, we first add environmental structure to a two-dimensional landscape of size 320×320 cells (see the Appendix section A2 for details). We do not define patches explicitly, but rather allow them to emerge from the spatial structure of the environment. We then seed *S* = 20 species onto the landscape, with initial biomass at each location drawn from a uniform distribution between 0 and 1, resulting in roughly equal biomasses at the landscape scale. For simplicity, we use periodic boundary conditions for the two-dimensional system (i.e., a torus topology), for both dispersal and interactions. We do not expect this choice to impact the results, due to the large size of the system considered.

We use 20 replicate landscapes, allowing environmental structure to vary among replicates while keeping the environmental scale constant. Replicates with other values of environmental scale are presented in the Appendix section A3. Each landscape replicate uses a different set of species and their interactions, chosen at random. Each replicate landscape was used to systematically vary the spatial scale of interactions *I* and dispersal coefficient *D*, with 25 different combinations (5 values of *D* and 5 values of *I*, as given in Fig. 2), giving a total of 500 simulations. We ascertain the generality of our findings by comparing across replicates.

**Figure 2:**
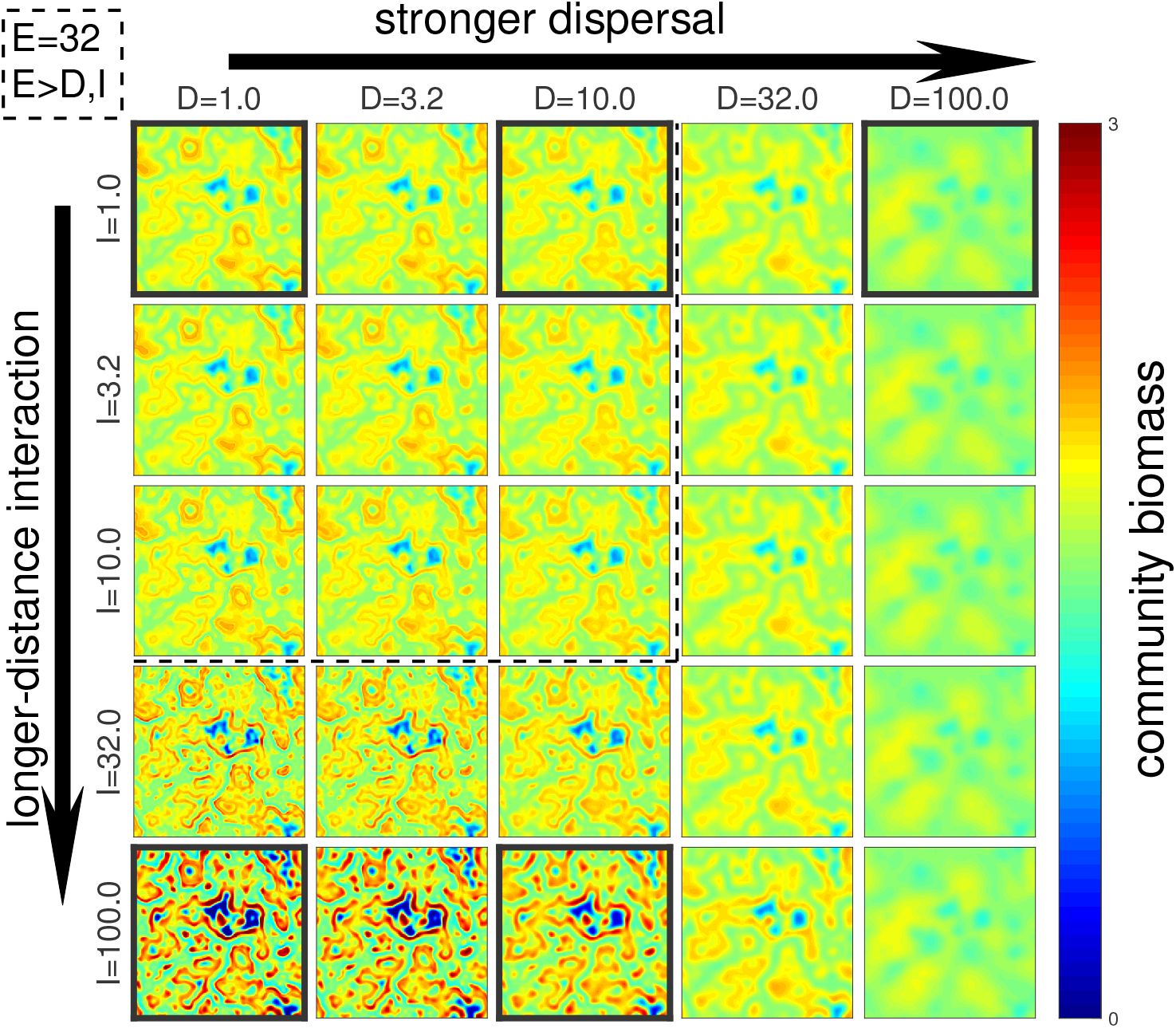
Distribution of total community biomass across the landscape as we change dispersal D (columns) and interaction I (rows) scales. Dashed black line shows where the environment scale E = 32 is larger than both D and I. Black frames around panels designate parameter values that we further examine in other figures. For better legibility, biomass levels above 3 are not shown.

We run each simulation, where a simulation is defined as a model run with a unique combination of process scales and replicate landscape, to a maximum time of *T* = 1000, or until equilibrium is reached. For practical purposes, we define an equilibrium as when the maximal change in biomass of any species in any location over a time-span of *T* = 1 is less than 10^−5^. A full list of parameter values can be found in Table 1. All simulations were performed using MatLab 2019a.

### Measurements

For each simulation we measure individual and total community biomass, species richness, and sample the landscape to calculate Species-Area Relationships (SAR curves) as well as Biodiversity-Ecosystem Functioning relationships (BEF curves). For species richness, SARs, and BEFs, we define a species to be extinct at a given location if its biomass is below than a threshold of 10^−3^.

To calculate SAR curves, we sample at 40 different spatial scales from 1×1 (single cells) to 320×320 (the entire landscape) on a logarithmic scale, and computed the species richness at each. For a given scale, we randomly choose 100 locations in the landscape, and sampled a region centered around the location chosen. We averaged over the 100 locations to obtain the mean richness value for a given scale.

We calculate both local and regional BEF curves, based on random sampling of the landscape. We do this in a similar way to the SAR curves, measuring species richness but also total community biomass. For the local BEF, we use a 1×1 cell area with 102,400 random locations chosen, while for the regional BEF we use an intermediate area of size 10×10 with 1024 locations sampled. In this way the BEF measurement is done consistently for different region sizes. For both local and regional BEF curves, we measure every cell on average once.

A striking outcome observed in our results is that spatial patterns of biodiversity and functioning in landscapes are not well captured by landscape summary measures, such as SARs. To explain these patterns, we calculate how correlated the biomass is of a given species as distance between sampling locations increases (i.e., spatial correlation), which can be used to quantify the properties of spatial patterns we observe. To calculate species’ spatial correlations, we do the following: 1) we normalize the species’ distribution by subtracting its average biomass (taken over the whole system); 2) we obtain a correlation map by calculating the convolution of a spatial distribution with itself, using a two-dimensional Fast Fourier Transform; 3) we normalize the correlation map by dividing the resulting two-dimensional map by its maximum value (i.e., we set a correlation value of 1 at the origin); and 4) we define the one-dimensional correlation function as the average between a vertical and horizontal transects through the correlation map. To define the scale of correlation for a given species, we locate the distance at which the correlation function reaches half its height, i.e., the distance from the origin where its value is the average of the maximum value (which is always 1) and its minimal value (typically around 0). A step-by-step illustration of calculating the spatial correlation is provided in the Appendix, Fig. S13.

## Results

### Local outcomes: functioning and diversity across localities

Our first major result is that, although they can arise from similar biological mechanisms (e.g., individual mobility), dispersal and interaction scales have opposite impacts on biodiversity and functioning patterns across the landscape (Fig. 2 and S9). We start from the case of weakly-connected communities with local interactions where all landscape patterns result from environmental variation (top-left panel, Fig. 2). Increasing the spatial scale of dispersal leads to a blurring of total community biomass over the landscape (from left to right, Fig. 2). In contrast, increasing the scale of species interactions leads to a sharpening of spatial patterns, amplifying underlying environmental heterogeneity (top to bottom, Fig. 2). The antagonism between these two effects can be seen by the fact that they counteract each other when increasing both scales at once, leading to similar-looking outcomes (along the diagonal, Fig. 2), but dispersal eventually wins out – the states along the right column are virtually identical, whereas the same is not true across the bottom row. Critically, it is not until the scales of dispersal or interactions exceed the scale of environmental heterogeneity (i.e., outside the dashed-lined boundary in Fig. 2) that the scale of either process significantly alters spatial patterns in biomass (see also Fig. S4). Larger emergent scales of total community biomass due to high *D*, and the opposite due to high *I*, can also be seen in Fig. 5, which shows how quickly patterns among locations become dissimilar as the distances between them increase.

We then focus on a subset of our scenarios above to show how process scales impact not only total biomass but also individual species distributions (Fig. 3). We observe that increasing dispersal scale predictably makes larger, more coherent domains (i.e., fairly defined areas with similar characteristics) with typically higher local diversity. Increasing interaction scale creates a more granular landscape with a broader range of diversities, including many low-diversity patches and a few high-diversity ones. Indeed, large interaction scales lead to more spotty species distributions, with rare species persisting in some locations where they would not in other scenarios (Fig. 3 bottom row). Two notable examples include species 1 (red patches) persisting only when interactions are large and dispersal is small, and species 2 (individually green, but here cyan due to its coexistence with species 3, blue) taking on a more clumped distribution with large interaction scales.

**Figure 3:**
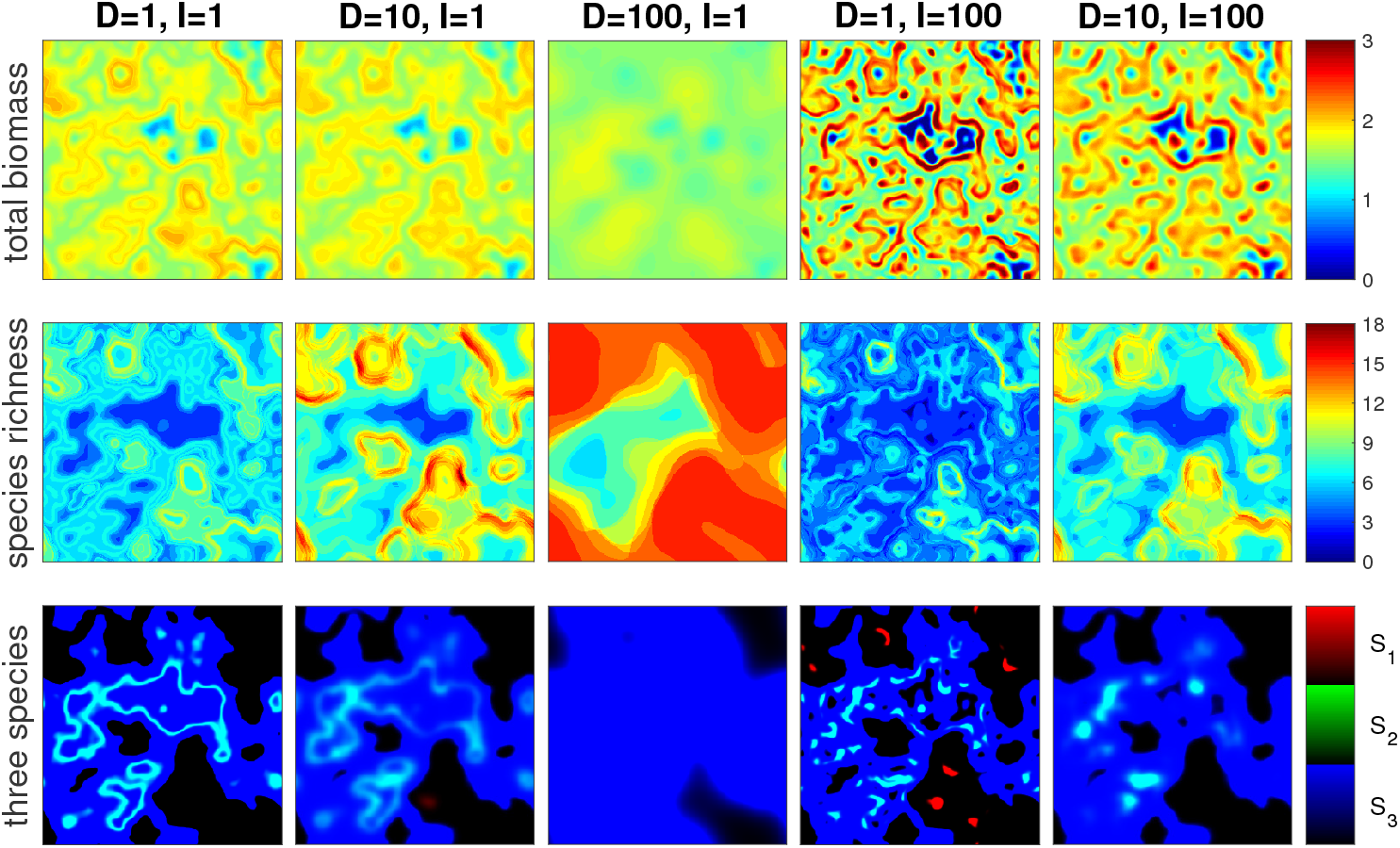
Species distribution patterns for five selected parameter sets, representing different scales of dispersal (D) and interaction (I), as designated in Fig. 2. **Top row**: total community biomass. **Middle row**: local species richness. **Bottom row**: distribution of three of the 20 species in original species pool (their biomass are encoded in the red, green and blue color channels, respectively; thus, cyan regions corresponds to coexistence of species 2 and 3). For better legibility, biomass levels above 3 are not shown.

### Regional outcomes: functioning and diversity at the landscape scale

The outcomes described above allow us to identify spatial patterns in local outcomes in the landscape, but what are outcomes for the landscape as a whole? Given the additive nature of biomass across localities, two regions could have identical biomass at the landscape scale even if one region has high variation among localities that span extremes of high and low values, whereas another varies little with biomass values that are intermediate. Here, we see that biomass is highest when interaction scales are large (Fig. S10), an effect that is quickly eroded as dispersal scales increase. Interestingly, these high-biomass landscapes had extreme variation in biomass among localities, including areas of extremely low biomass (dark blue in Fig. 2) and extremely high biomass (red in Fig. 2). Therefore, high biomass is driven by a disproportionate subset of local communities in a landscape. Furthermore, these high biomass landscapes were unremarkable in regional species richness in the landscape and actually had fewer species per locality on average than other scenarios (Fig. S11). For those who may be interested in comparing our findings to those typically reported in traditional metacommunity models more explicitly (e.g., [43], we note that the left and right plots in Fig. S11 essentially show local (i.e., alpha) and regional (i.e., gamma) diversity, respectively, whereas compositional turnover among localities (i.e., beta diversity) is essentially differences between them.

### Cross-scale outcomes: BEF and SAR

Next, we turn to two types of cross-scale outcomes (Fig. 4). First, we consider the relationship in BEF curves (i.e., total biomass vs. species diversity) at neighborhood (i.e., single cell) scales. In doing so, we find that BEF curves (Fig. 4, left panel) reflect underlying process scales. In particular, they exhibit a hump-shaped relationship for large interaction scales, suggesting that patches with the largest total biomass are not the most diverse, but rather have a few high-performing species. This result ties into our previous observation that the interaction scale tends to amplify environmental heterogeneity, and may thus put more weight on selection effects, where abiotic conditions select the best-performing species at the exclusion of others. We also examined BEF curves measured at larger scales, i.e., when spatially aggregating 100-cell neighborhoods, and found qualitatively identical patterns (Fig. 4, middle panel).

**Figure 4:**
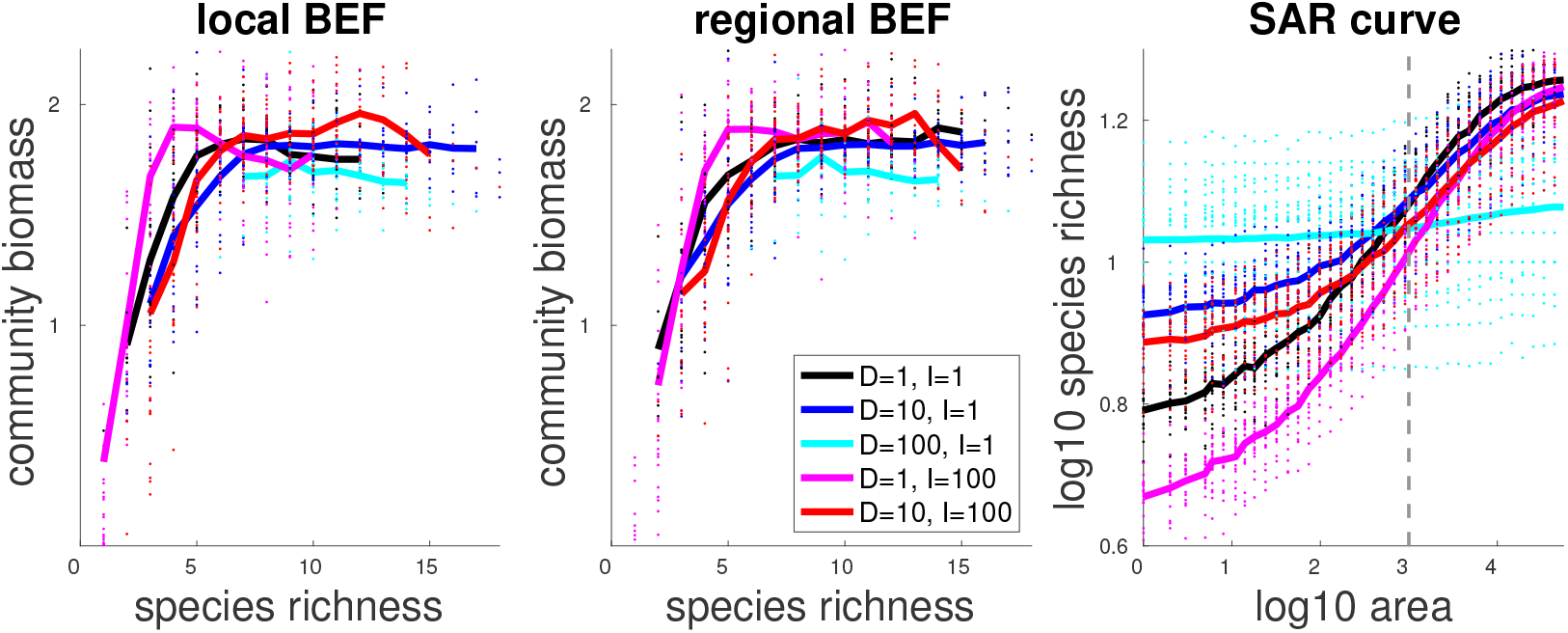
BEF and SAR relationships. . Solid lines show average values over 20 replicates, small circles show values for individual replicates. Colors correspond to five selected parameter sets, representing different scales of dispersal (D) and interaction (I), as designated in Fig. 2. Local and regional BEF curves are measured at regions of size 1 and 100, respectively. Vertical gray line shows the area corresponding to the environmental scale E = 32. Although our model is deterministic (i.e., each replicate has only one possible outcome, given a specific set of parameter values and initial conditions), differences among replicates reflect differences in parameter values caused by sampling those values from distributions (Table 1).

We also look at a pattern aggregated over continuously increasing spatial scales – the SAR (Fig. 4, right panel). We would expect that changes in the slope or shape of the SAR as the aggregation scale (x-axis) exceeds the spatial scales of our ecological processes, as has been demonstrated for the Stability-Area Relationships [8]. However, we do not observe a clear link between process and pattern scales, beyond the fact that the inflection point (in particular, for low *D* and *I*) corresponds to the environmental scale *E* (vertical gray line in Fig. 3). The main impact of process scale is that, by amplifying landscape heterogeneity, a large interaction scale *I* leads to a stronger SAR when large interaction scales are coupled with short dispersal scales. Specifically, as predicted, at the smallest scale the *D < E < I* scenario (magenta curve) yields the lowest species richness compared to all other scenarios, whereas at the scale of the entire landscape, its richness is very high.

Aggregated measures of biodiversity and functioning at regional scales miss much of the information captured by local measures, such as the distribution and turnover in biomass (Fig. 2 and Fig. 3). Yet these local patterns can be quantified. Figure 5 presents the results of the spatial correlation of species biomass distributions, which measures how the biomass of a species correlates over the distance between sampling. We observe clear trends in scale, with consistent patterns of growing (shrinking) correlation with higher dispersal (interaction) scales.

**Figure 5:**
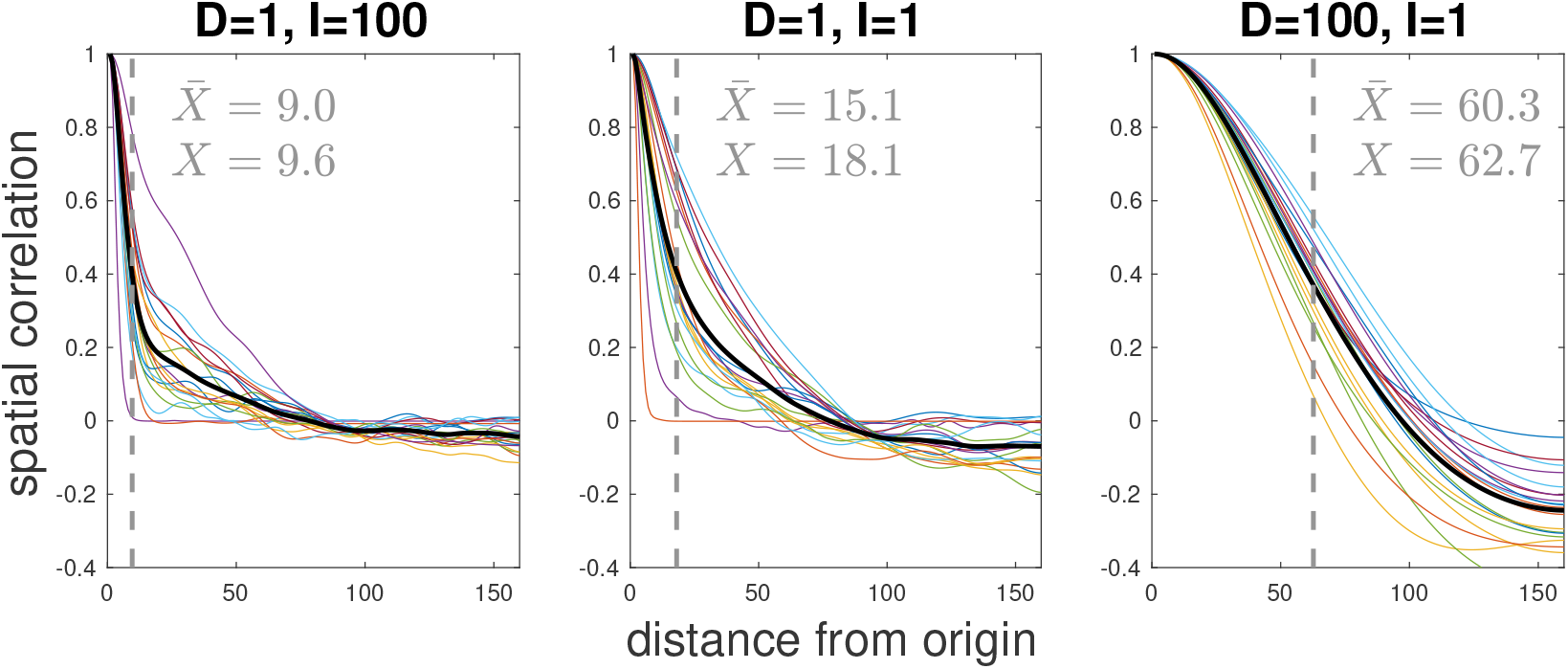
Spatial correlation of each species’s biomass distribution, for three scenarios. **Left**: I = 100, D = 1; **Middle**: I = 1, D = 1; **Right**: I = 1, D = 100. Recall that E = 32. Each of the 20 species is represented by a different color, with black showing the average correlation function, all for a single replicate. For this simulation run, the scale of correlation X is given, and is shown by gray vertical lines. The correlation scale averaged over the 20 replicates, 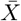, is also noted.

## Discussion

This study focuses on a critical question: how is the scaling of ecological patterns, such as patterns of biodiversity and ecosystem functioning, related to scales of specific processes, and why? We have modelled how intrinsic scales of ecological processes align with the emergence of ecological patterns in a metacommunity, where we control the spatial scale of environmental heterogeneity, dispersal, and species interactions. In doing so, below, we highlight the following three take-home messages of our results:

- the scale of one process (here, environment) can cause the emergence of characteristic scales of other processes (dispersal, interactions)
- two interlinked ecological patterns (biodiversity and ecosystem function) and their relationship to each other are oppositely affected by two forms of organismal movement
- averaging ecological patterns at any one scale misses a rich patterning of spatial variance that is closely tied to process scales

Below, we expand upon each finding and place them within existing knowledge, examine the mechanisms that underlie our findings, contrast results among ecological variables, and end by placing our results within a context of ecosystem preservation.

A main finding of our study is that the spatial scale of interactions amplifies environmental heterogeneity, sharpening observed spatial patterns, in contrast to dispersal scales. Importantly, observed spatial patterns did not reflect the absolute value of the spatial scale of each ecological process, but rather, their values relative to the environment; decreasing the spatial scale of the environment shifts the boundary of blurring/sharpening effects of dispersal and species interactions (Fig. S4). We find this effect because environmental conditions are quite literally the template upon which dispersal and species interactions mold species distribution. Large-scale (i.e., at scales above the template) processes are more important than small-scale ones in determining overall patterns, meaning that only when dispersal or interactions have large scales can they impact large-scale patterns.

We examined the impacts of process scales on two classes of patterns: first, on the spatial scaling of patterns (SAR and BEF), and second, on the spatial structure of species biomass in the landscape. Unexpectedly, the latter class of patterns appears to better reflect the scale of ecological processes, such as the distribution and turnover of biomass and biodiversity across the landscape. These patterns would be lost by examining mean biodiversity and function at specific aggregation scales (e.g., local vs. regional; Fig. S4), but were well captured via spatial autocorrelation (Fig. 5). From these analyses, one take-home message is that increasing the scale of species interactions actually amplifies variation on small scales. In other words, large-scale processes do not necessarily beget large-scale patterns.

The question of how process scales affect observed patterns can also be spun around: what information about process scales can be inferred from the various patterns we see? Considering the opposing effects that dispersal and interaction scales have on pattern scales (Fig.2), it is not clear that such an inference is possible. However, given that patterns scales change differently (compare Fig. 2 with Fig. S3, for instance), combining several measures together may provide an answer, for instance by finding when changes in spatial correlations of biodiversity and biomass no longer behave similarly. In this context, it is perhaps to be expected that no clear connection was found between well known patterns such as BEF and SARs, and process scales. Over the past few decades, ecologists have been cautioned from interring processes from patterns [44]. Our results demonstrate exactly why this is important: a lack of a 1:1 mapping between a pattern and any one specific process.

Indeed, our finding that the SAR curves did not exhibit transitions at particular spatial scales, that would allow us to identify the typical scales of the underlying processes (other than the environment), runs counter to other contexts, such as the invariability-area relationship [8]. In particular, we do not find a triphasic SAR curve that is often reported [45, 8]. This is the case since our model does not consider individual sampling and dispersal limitation, which typically lead to stronger SAR slopes at small and large scales, respectively. We thus see the strongest slopes at intermediate spatial scales, consistent with results under similar settings [46], and hinting that we are largely seeing community dynamics typical of species-sorting [37]. Centering on the average SAR slope itself, on the one hand, we found that large interaction scales may enhance the SAR by amplifying landscape heterogeneity and creating low-diversity strips along the edges of species ranges. On the other hand, this spatial heterogeneity could also promote coexistence as a weaker competitor might thrive in the margins [47]. This suggests that edge effects may play a prevalent role in the case of long-range interactions, and deserves more extensive investigation. Overall, the scales of biotic processes (interaction and dispersal) are mainly reflected inasmuch as they change overall community properties, such as total diversity across the landscape.

Knowledge of the spatial scale of ecological processes is critical to understanding the maintenance of ecosystems. To illustrate this argument, one can imagine a landscape manager interested in preserving some baseline level of functioning in a landscape at a specific spatial extent, for example, primary production. If the spatial scale of interest does not encompass the intrinsic scales of processes that govern functioning, then landscape alteration beyond that scale might impact functioning in an unanticipated and undesirable manner; these scales will differ among ecosystems based on how species? traits and the physical landscape affect how organisms experience scales of E, D, and I. In other words, the scales important to the maintenance of ecosystem function may be mismatched from the (typically small) spatial scales at which ecosystem functioning is observed and managed, but the degree to which this is true depends on process scaling. Predictions of our model could be best tested empirically in microcosm or mesocosm setups or using data syntheses, for example, by examining the spatial structure of species richness and biomass depending on process scales of focal taxa (e.g., small vs large-bodied animals using remotely sensed data, experiments with insects where mobility is restricted).

Our results suggest that it will be difficult to manage landscapes to preserve biodiversity and ecosystem functioning simultaneously, despite their causative relationship, for two related reasons. First, the fact that increasing dispersal and interaction scales had opposing effects on either ecosystem property presents a unique management challenge, given that both scales are tied to organismal movement, albeit on distinct timescales (i.e., daily vs. once-per-generation). Second, ecosystems attained the highest biomass in scenarios which also led to the lowest levels of biodiversity, specifically, when interaction scales were large and dispersal scales were small. We note that this second issue may only be relevant when interactions are largely competitive, since our modeling, and thus results, did not consider mutualistic interactions which would likely change the observed trade-off between biodiversity and biomass. How would a manager plan a landscape to enhance interaction scales (preserving function) while simultaneously minimizing scales of dispersal (preserving biodiversity)? This can, for instance, be relevant for managing predation of pest herbivores in agricultural landscapes [16]. This type of intervention might be most successful in species with body plans for long-distance movement, but that can remain philopatric for behavioural reasons (which can be environmentally determined; i.e., territorial hunters).

Our metacommunity model differs from traditional metacommunity models in several important ways. Traditional metacommunity models tend to include discrete local patches embedded within an implicit inhospitable matrix, interconnected by rates of dispersal, often from a spatially-implicit regional pool of dispersers. By contrast, “patches” in our model emerge from the environmental template (Fig. 3), the structure of which may be viewed differently by different species according to their fundamental niche. Further, these patches may have fuzzy boundaries, within-patch heterogeneity, as well as different shapes and sizes. Individuals may be lost to the matrix (i.e., habitat falling outside of the fundamental niche) if they disperse there or may form stepping stone populations to reach new patches. In doing so, dispersal limitation is more likely to emerge as the spatial grain of the environment exceeds the scales at which species disperse, a major result of our study. These features align with the recent calls [48, 31] to develop more realistic metacommunity models applicable to a wider range of systems, beyond discrete, patchy, island-like systems. Given these strengths, the next step is to extend a model like ours to multi-trophic systems, beyond “horizontal” (sensu Vellend [49]) competitive communities. Our model is naturally amenable to multitrophic systems, as predators often perceive the landscape at a different scale than their prey (i.e., a different interaction scale) and would perceive the scale of the environment via spatial distributions of their prey–additionally, there is an opportunity to move beyond Lotka-Volterra dynamics for modelling species interactions, towards more mechanistic consumer-resource approaches [50]. Most metacommunity models have been applied to competing species [17], with multi-trophic extensions becoming more common in recent years [51].

Our conclusions are twofold. First, we bring forward an important spatial scale – the range of species interactions – that has been largely neglected in previous analyses (e.g., metacommunity theory). This interaction range can be derived from many of the same ecological mechanisms as dispersal, such as individual mobility, yet these two processes lead to opposite ecological effects. This suggests that we must carefully distinguish whether mobility actually leads to population dispersal or to large-range interactions, and re-evaluate possible consequences of evolution or environmental change in these processes. Finally, we saw that the spatial scale of ecological processes might not appear clearly in the scale of resulting patterns such as Species-Area or Biodiversity-Ecosystem Functioning relationships, though they may sometimes be reflected in local outcomes. While we focused on a few important biodiversity and functioning patterns, our study paves the way for future work investigating systematically under which conditions various ecological pattern scales may or may not reflect the spatial scale of underlying processes.

## Acknowledgements

YRZ, MB, DWS and ML were supported by the TULIP Laboratory of Excellence (ANR-10-LABX-41), and by the BIOSTASES Advanced Grant, funded by the European Research Council under the European Union’s Horizon 2020 research and innovation programme (grant agreement no. 666971). We thank all members of the BEF Scale working group for valuable discussions and feedback.

## Data accessibility

Script files for simulations and analysis of results shown in the manuscript are available at the open-access repository: https://doi.org/10.5281/zenodo.5543190.

## Appendix

This appendix is made of four sections. A1: Measurement of scales; A2: Generating the landscape; A3: Different environmental scales; A4: Additional plots.

### A1 Measurement of scales

As explained in the main Methods section, we explicitly measure and compare three spatial scales: environmental conditions (E), dispersal (D) and species interaction (I). We now detail the definition of these three scales, and finally note the peculiarity of dispersal scale.

#### Environmental scale E

The environment itself is generated using a combination of a spectral color and cutoff wavenumber (see next section), but this does not explicitly define the scale. To measure the scale of the environment, we follow the same approach as for the correlation function and measure the scale for a species biomass distribution (using a convolution based on FFT), except that we do this for the value of intrinsic growth rate 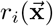, as it is directly set by the environment. For each of the 20 species, we can calculate a correlation function (in the same manner as explained in the methods), and from this we calculate the correlation scale (the point of middle height for the correlation function). We average this value over all 20 species, to calculate the environment’s scale for a given system. Since this result depend on the randomization of the environment, we repeat this for many replicates, and choose values of *ρ* and *k*_*c*_ that will on average give a value of *E* we want to have.

#### Dispersal scale D

To estimate the dispersal scale *D*, we compare the effect of changing the dispersal coefficient *δ* with changing *γ*. In Fig. S1 we show how changing *δ* and *γ* (and thereby *D* and *I*) affects the community biomass distribution. As seen in the left panel, with low *δ* and *γ* the difference from a null scenario of no dispersal and no interaction distance is very small, but increasing either *δ* or *γ* changes the community biomass distribution considerably. In the middle and right panels we see these differences, as we change only *δ* (middle) or only *γ* (right). This clearly shows three things: 1) The effect of interaction distance scales linearly <u>w</u>ith *γ*, as expected by its definition. 2) The effect of dispersal coefficient scales with 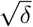, as expected from dimensional considerations (e.g., [42]). 3) More specifically, to ma<u>k</u>e these two effects comparable, the dispersal scale is missing a factor of 10, i.e., 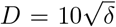. This can be seen by the fact that for both *δ* = 1 in the middle panel and *γ* = 10 in the right panel, the y-axis values are roughly the same (10^−1.2^).

#### Interaction scal I

In our model, the species interactions are explicitly defined with a distance over which they occur – via the Gaussian kernel function. This naturally gives us the scale of interactions *I*, as the width of the Gaussian function, such that *I* = *γ*.

**Figure S1:**
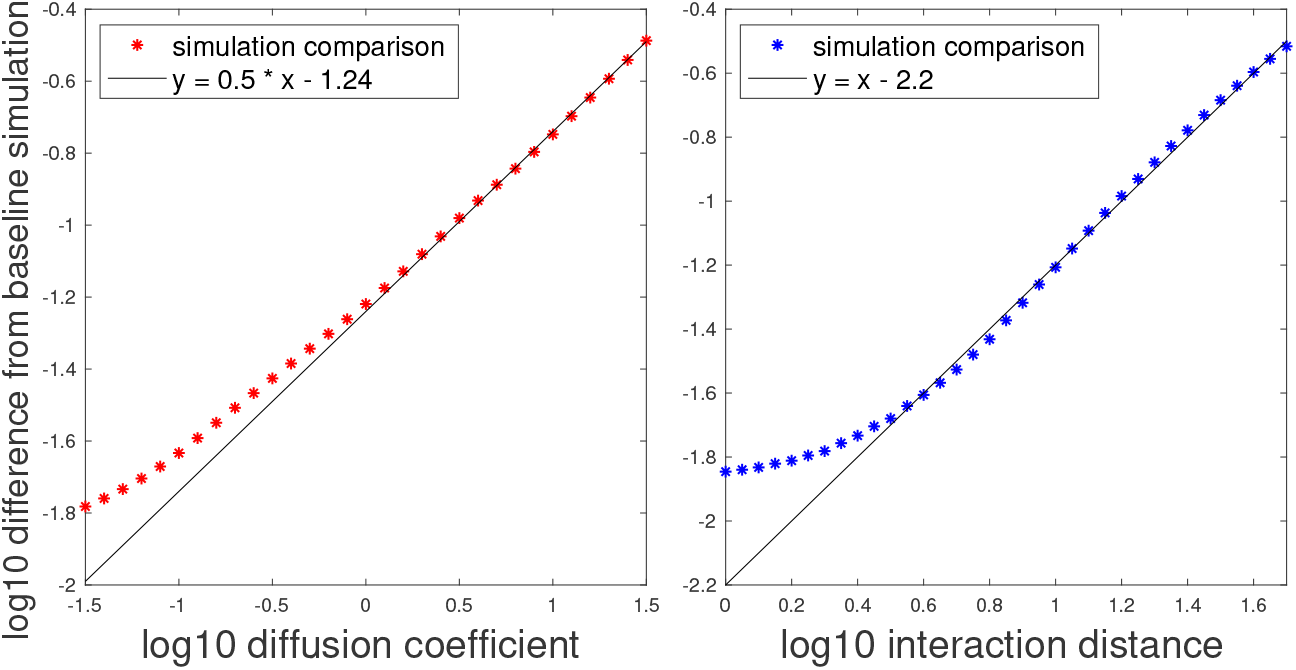
Comparison of different diffusion coefficient and interaction distance scenarios to the case of no dispersal and local interactions alone. Differences are squared, summed over all species, and averaged over domain. This is done along the diffusion coefficient (interaction distance) axis in the left (right) panel. Comparison shows that diffusion scales like a square root, and that a normalization factor of 10 should be applied to make it comparable to interaction distance (i.e., d = 1 is comparable to an interaction distance of 10).

#### Peculiarity of dispersal scale

An interesting problem we encountered, which is worth expounding upon to aid future research in this area, is how to place dispersal on comparable scales and strength to other processes. For both environmental factors and species interactions, we could separate the intensity of variation and the scale over which it takes place. We could do this, for instance, by modelling interactions with a spatial kernel which defines the range of these interactions. For dispersal, however, this distinction does not hold in the continuum limit nor in the stable equilibrium regime that we consider in this study. This can be understood intuitively in a single dimension: organisms who disperse from site *x* to site *x* + 1 at time *t* will be counted in those that disperse from site *x* + 1 to site *x* + 2 at a later moment in time. Therefore, dispersing twice as fast between neighboring sites can be equivalent to dispersing twice as far. This equivalence breaks down when the details of individual dispersal events matter, e.g., for very rare and long-ranged dispersal events [52]. But even then, the strength of each dispersal event would still play into the spatial scale over which dispersal impacts the dynamics over longer times. As a consequence, defining dispersal scale from a spatial kernel alone might seem more intuitive, but would actually hide the importance of intensity, and we prefer to simply model nearest-neighbor dispersal and acknowledge that intensity and scale are entangled.

### A2 Generating the landscape

The landscape profile is defined by a spectral color (*ρ*) and cutoff (*k*_*c*_). A spectral color close to 0 corresponds to “white” noise, i.e., noise that exhibits little or no spatial autocorrelation; a spectral color close to 1 indicates “red” noise – noise with high spatial autocorrelation [40]. The spectral cutoff creates a point of truncation in the frequency profile that prevents high variation between adjacent cells, in effect smoothing the noise across the landscape. Together, color and cutoff control the degree of structural fragmentation of the landscape (see Fig. S3). More weight on higher frequencies (low *ρ*, high *k*_*c*_) entails smaller and less-connected fragments of similar environmental conditions. Weight on lower frequencies (high *ρ*, low *k*_*c*_) creates long bands of constant environmental conditions which can act as corridors for species favoring this value.

To generate the environmental landscape 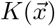, we prescribe a frequency profile for the noise:

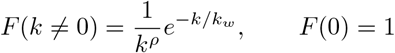

which is a power-law with color *ρ* (*ρ* = 1 corresponds to red noise) and an exponential cutoff with wavenumber *k*_*w*_ = *k*_*c*_*L/*2 which removes high spatial frequencies, smoothing the landscape and avoiding strong variations between adjacent cells. The construction process is demonstrated in Fig. S2. Note that the cutoff wavenumber is simply the normalization of the spectral cutoff by the number of different frequencies represented by the chosen resolution of the domain, *L/*2, with *L* the number of cells along the x and y axes, such that in the spectral domain it represents the resolution of the landscape.

**Figure S2:**
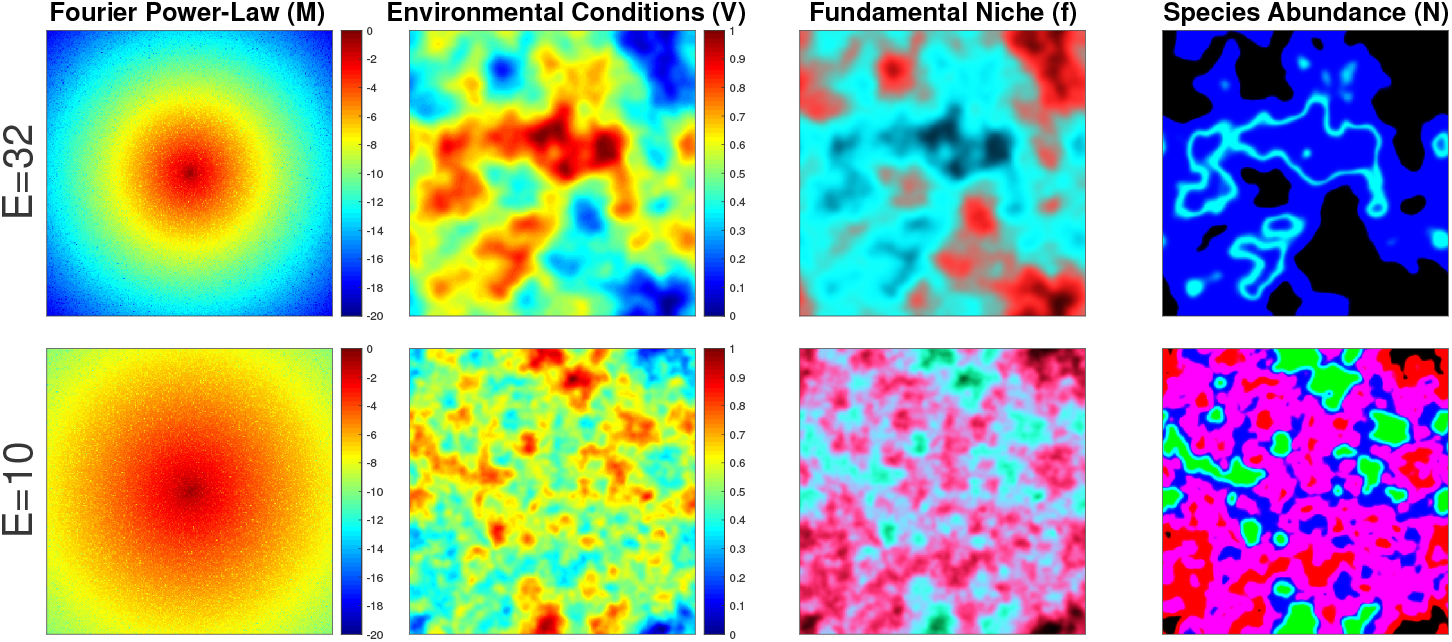
Demonstration of landscape construction. The steps of landscape construction are shown in the different columns, with the top (bottom) row corresponding to a landscape with E = 32 (E = 10). From left to right, the four columns correspond to: 1) The function M, which is a power-law function with exponential cutoff, on a two-dimensional spectral map (i.e., where each cell corresponds to a different spatial frequency), with the addition of random noise. 2) The environmental conditions V, which result from applying the Fourier transform on the previous step, and normalizing the values to range between 0 and 1. 3) The fundamental niches f_i_ of 3 species, where the value of f_i_ of each species are encoded in the red, green and blue color channels. 4) The spatial distribution of species biomass N_i_ at equilibrium, of the same 3 species and with the same color coding, as the previous column. Note that the top-right panel corresponds to the bottom-left column of Fig. 3.

Practically speaking, for a two-dimensional landscape, we generate a *L × L* matrix *R*_*ij*_ of uniform random numbers over [−1, 1] corresponding to amplitudes for each wave vector (*k*_*x*_, *k*_*y*_). We then multiply these random numbers by the profile above

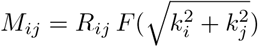

with 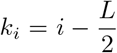 where index *i* is a natural number running over [1,L]. We set the element *M*_*L/*2,*L/*2_ corresponding to the uniform trend (*k*_*i*_ = *k*_*j*_ = 0) to 5. Finally, we apply a Fast Fourier Transform on the matrix *M*_*ij*_ to obtain the landscape matrix *V*. As explained in the main text, this landscape matrix *V* is used to define the growth rate *r*_*i*_ using a Gaussian function (see eq. 2), which in turn determines the species biomass distribution *N*_*i*_ (see eq. 1). We show in Fig. S3 the environment as a function of different values of *ρ* and *k*_*c*_, to better visualize their effect.

**Figure S3:**
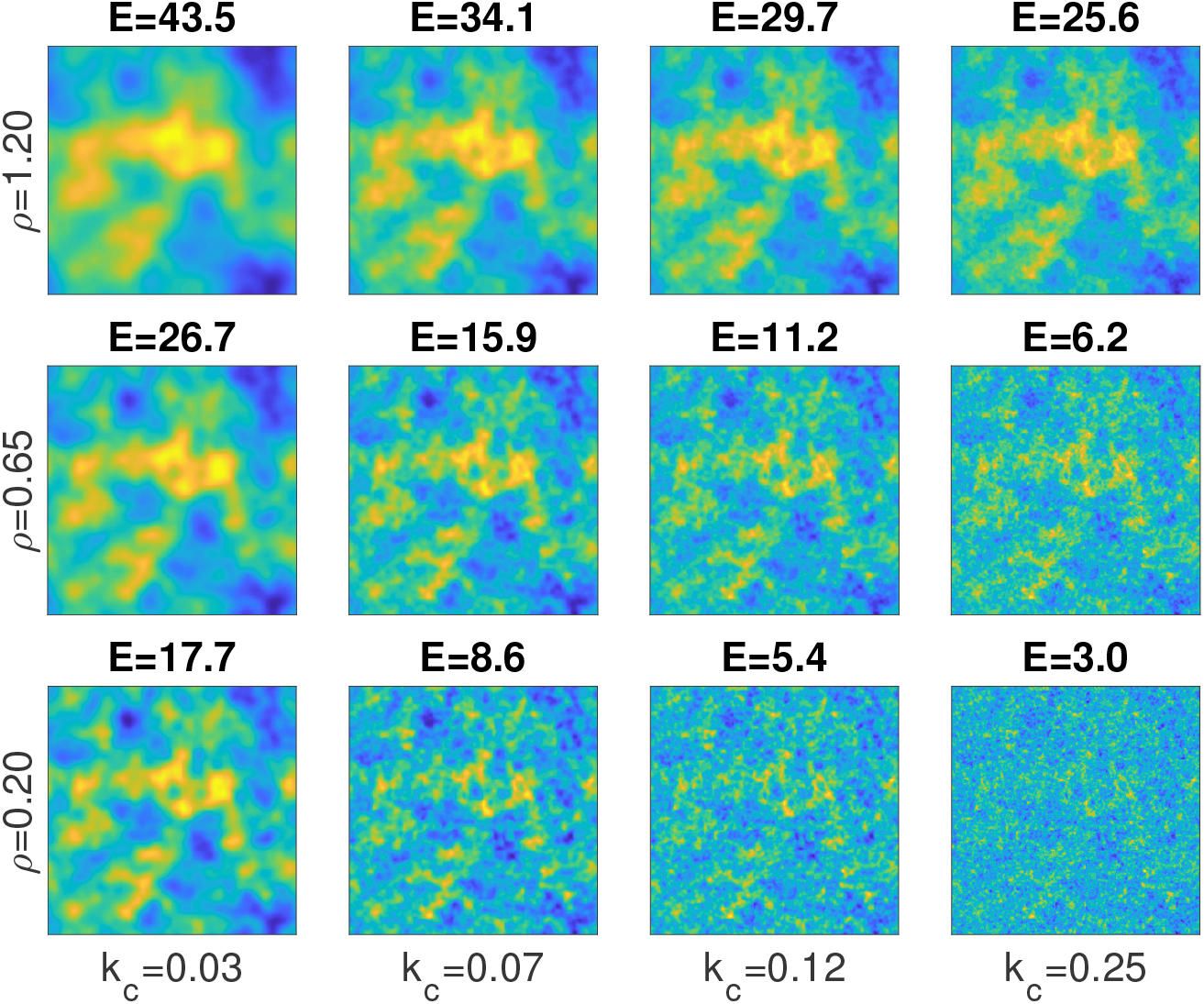
How ρ and k_c_ shape the landscape structure, shown by maps of the environmental conditions V. We show an example of how a landscape is affected by different values of ρ (rows) and k_c_ (columns). On top of each panel we also note the environmental scale E that corresponds to the combination of ρ and k_c_. We can see that smaller k_c_ values lead to a landscape with less sharp transitions (i.e., smoother), whereas ρ has a more significant effect on the overall scale. In other figures and in the main text we choose ρ and k_c_ concordantly, with large ρ values together with small k_c_ values for a large E, and small ρ values together with large k_c_ values for a small E.

### A3 Different environmental scales

We show below a few additional plots, which explore the impact of different values of environmental scale *E*. In Fig. S4 we show the overall difference in community state, between different sets of values of *D* and *I* to the case of no dispersal and local interactions, for two values of *E*.

**Figure S4:**
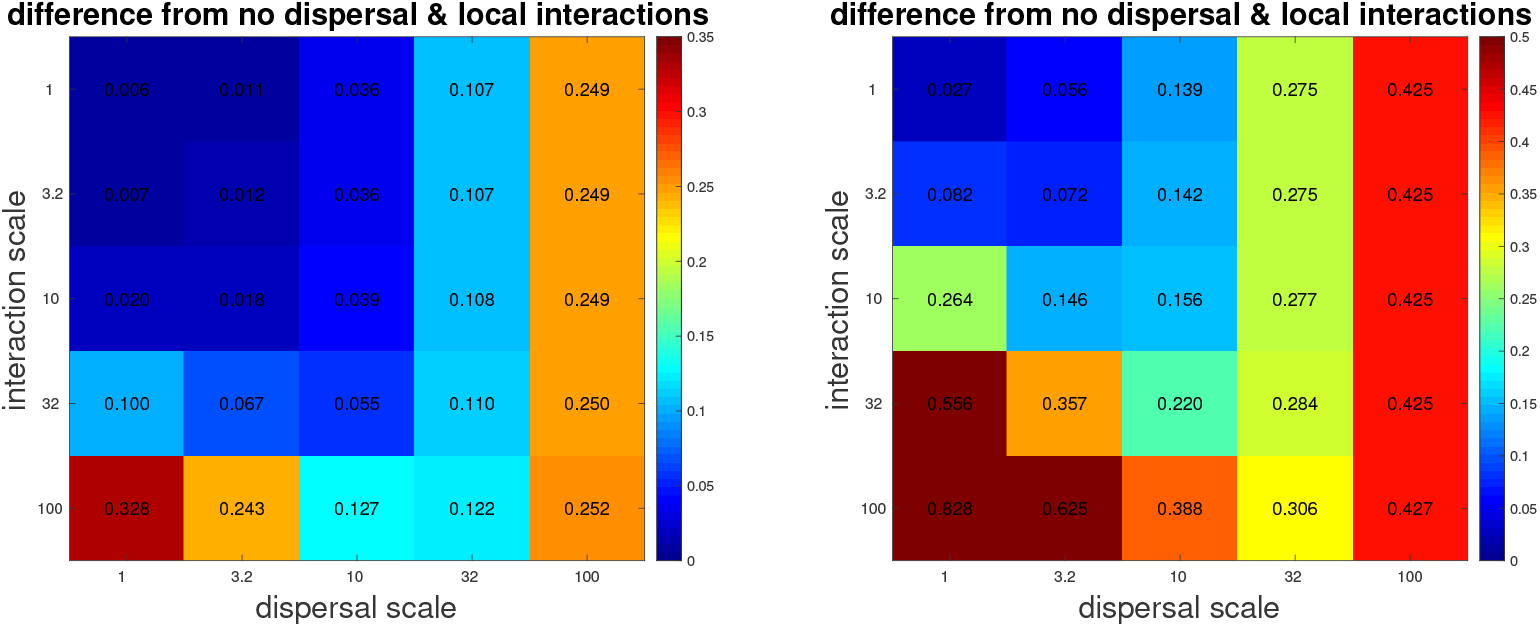
Comparison of various scenarios to the case of no dispersal and local interactions alone. Difference is measured by averaging over the squared sum of each cell for a given value of I and D, against the baseline of D = I = 0. This is done for for 5×5 different parameter sets with different values of D and I, for two different values of E, 32 and 10, in the left and right panels, respectively.

In Fig. S5 we consider different *E* values, and see how changing either *I* or *D* affects the overall change in system state (compared with the baseline of no dispersal and local interactions). In both figures we can see that big differences in the state of the system due to higher *I* or *D* (seen as dark blue region in Fig. S4, and region below the dotted line in Fig. S5) occur for lower values for *I* and *D*, and only when *E* is sufficiently high. This demonstrates that the environmental scale *E* determines the threshold scale of *I* and *D* in which they can have a substantial effect on the community.

We also test how the inflection point of SAR (measured in the same way as in the main text), changes along a range of *E* values (Fig. S6). We can see that as long as dispersal is not too high (i.e., the three cases where *D* = 1), the inflection point follows the environmental scale *E* (seen by the roughly parallel lines to the 1:1 line).

In Fig. S7 and Fig. S8 we show the spatial distributions of biomass and species richness, for a different landscape, one that has an environmental scale of *E* = 10.

**Figure S5:**
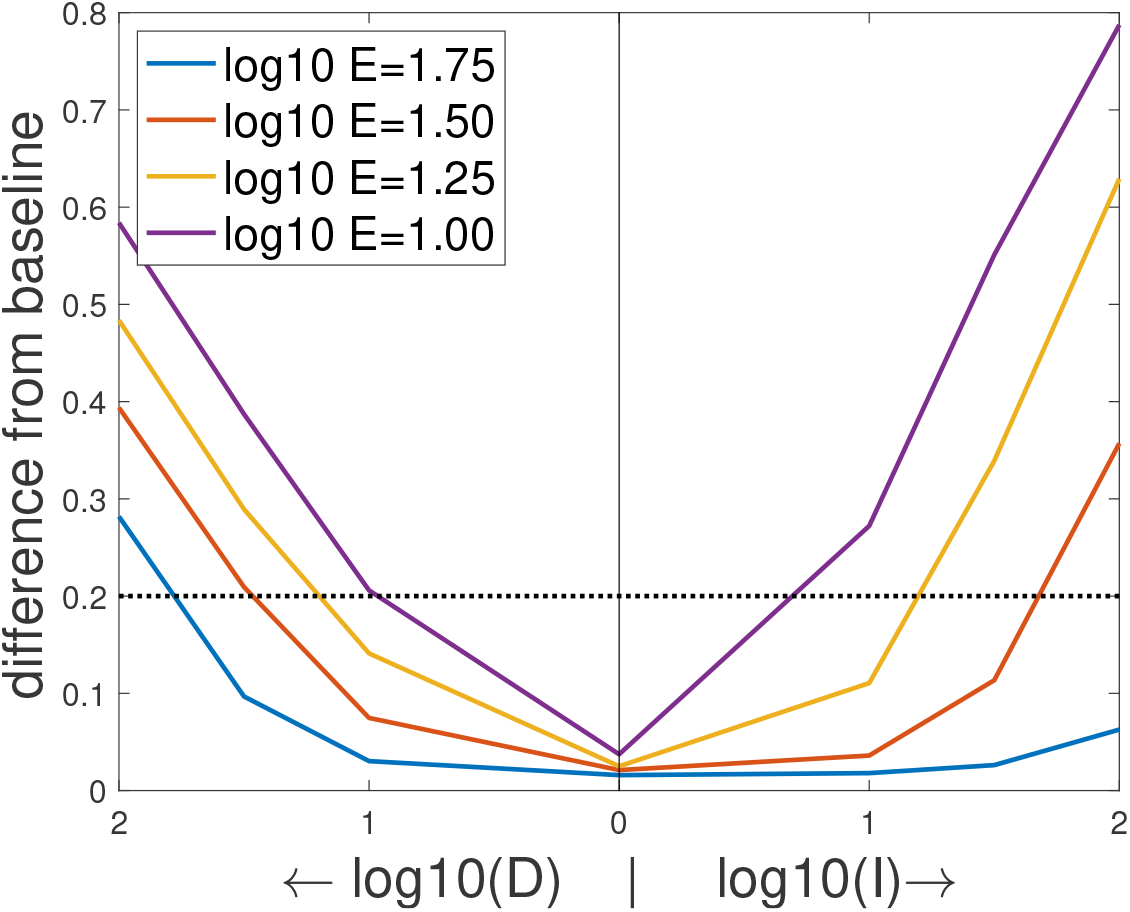
Average difference from a community with no dispersal and local interactions only. Difference is measured by averaging over the squared sum of each cell for a given value of I and D, against the baseline of D = I = 0. Left half shows the effect of D alone, while right half shows the effect of I alone.

**Figure S6:**
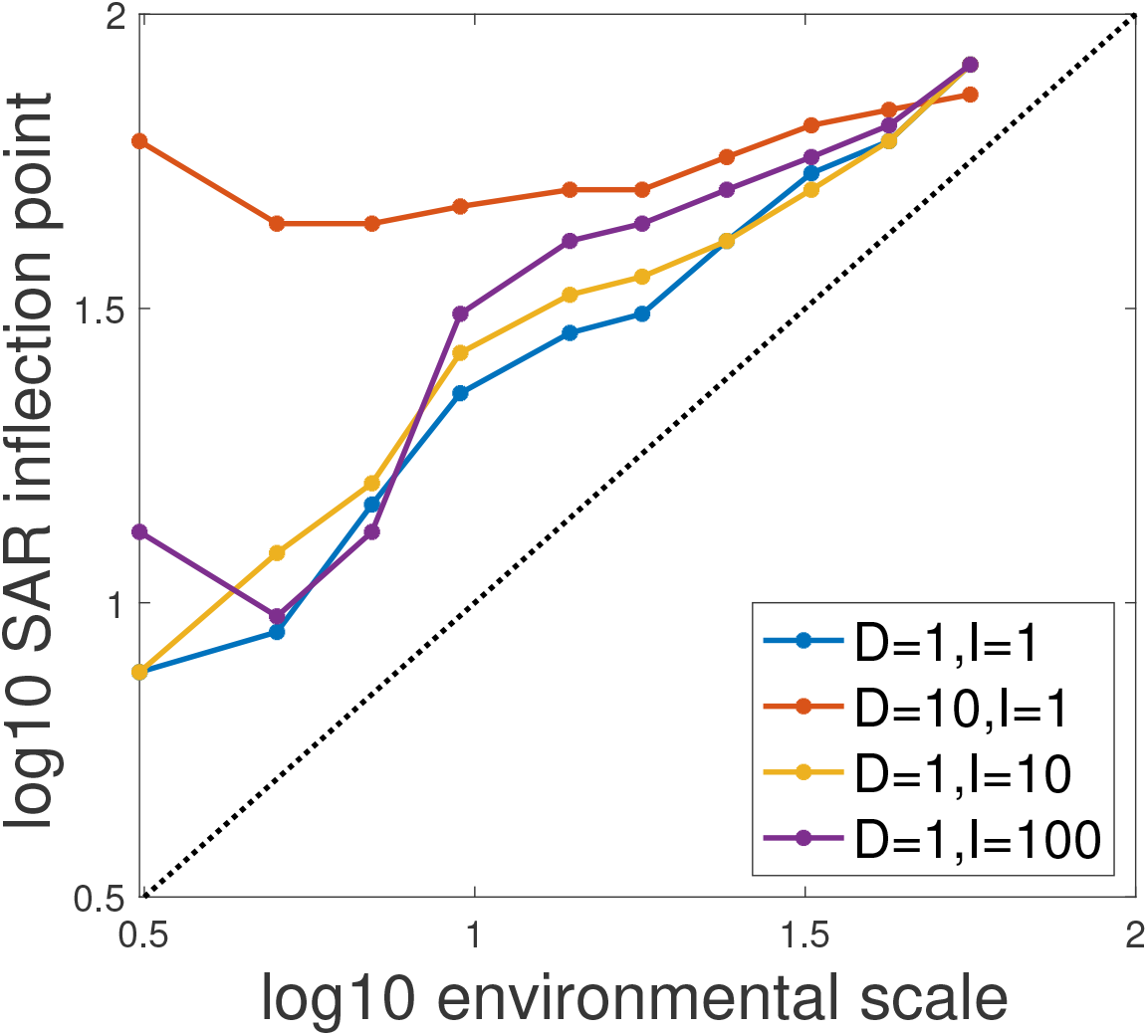
Inflection point of SAR for different combination of scales. For four sets of values of I and D (D = 1, I = 1 ; D = 1, I = 10 ; D = 1, I = 100 ; D = 10, I = 1), we show how the inflection point of SAR changes along a range of 10 values of E (with values between 56 and 3).

**Figure S7:**
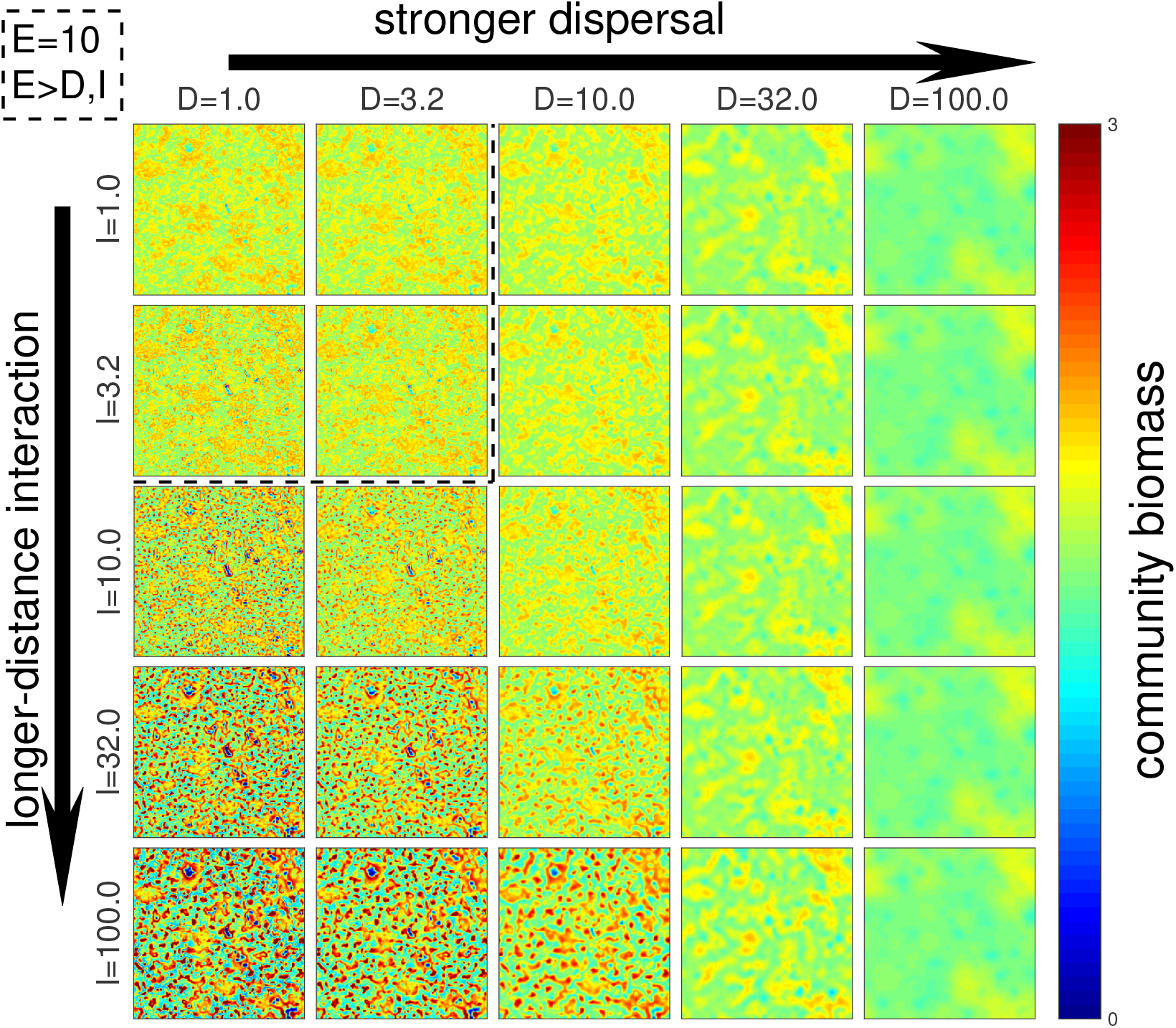
Total community biomass, for the 5×5 scenarios, with E = 10. For better legibility, biomass levels above 3.0 are not shown.

**Figure S8:**
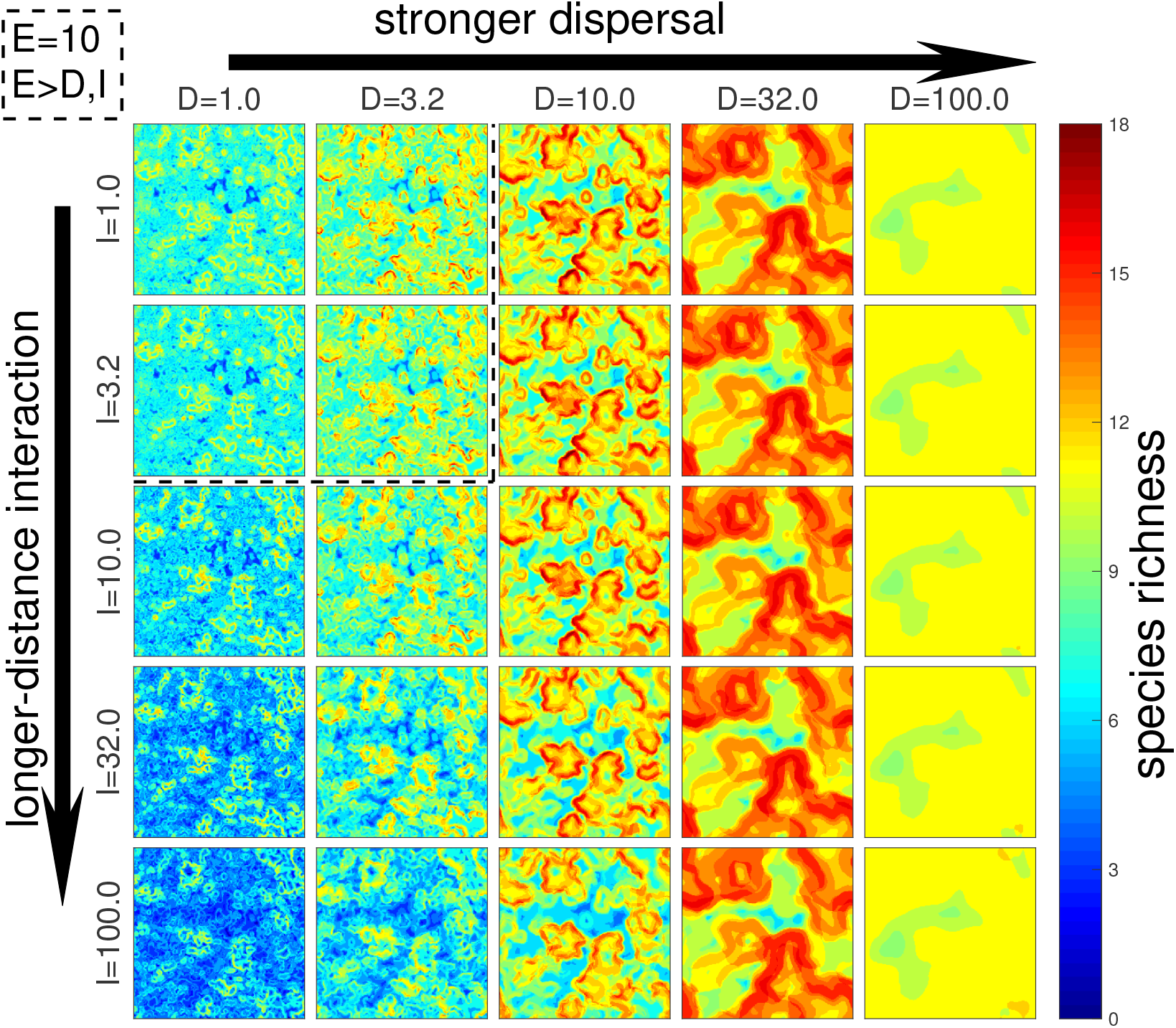
Local species richness, for the 5×5 scenarios, with E = 10.

### A4 Additional plots

We show below a few additional plots.

In Fig. S9 we show the spatial distribution of species richness, for 5×5 different parameter sets with different values of *D* and *I*, corresponding to Fig. 2. In Fig. S10 and Fig. S11 we show summary statistics for each of these 5×5 parameter sets, of total community biomass, average local diversity, and total diversity.

Finally, we explore in Fig. S12 the sensitivity of our results to the parameter *β*, and demonstrate using Fig. S13 the calculation of species’ spatial correlations, which is used to estimate the environmental scale *E*.

**Figure S9:**
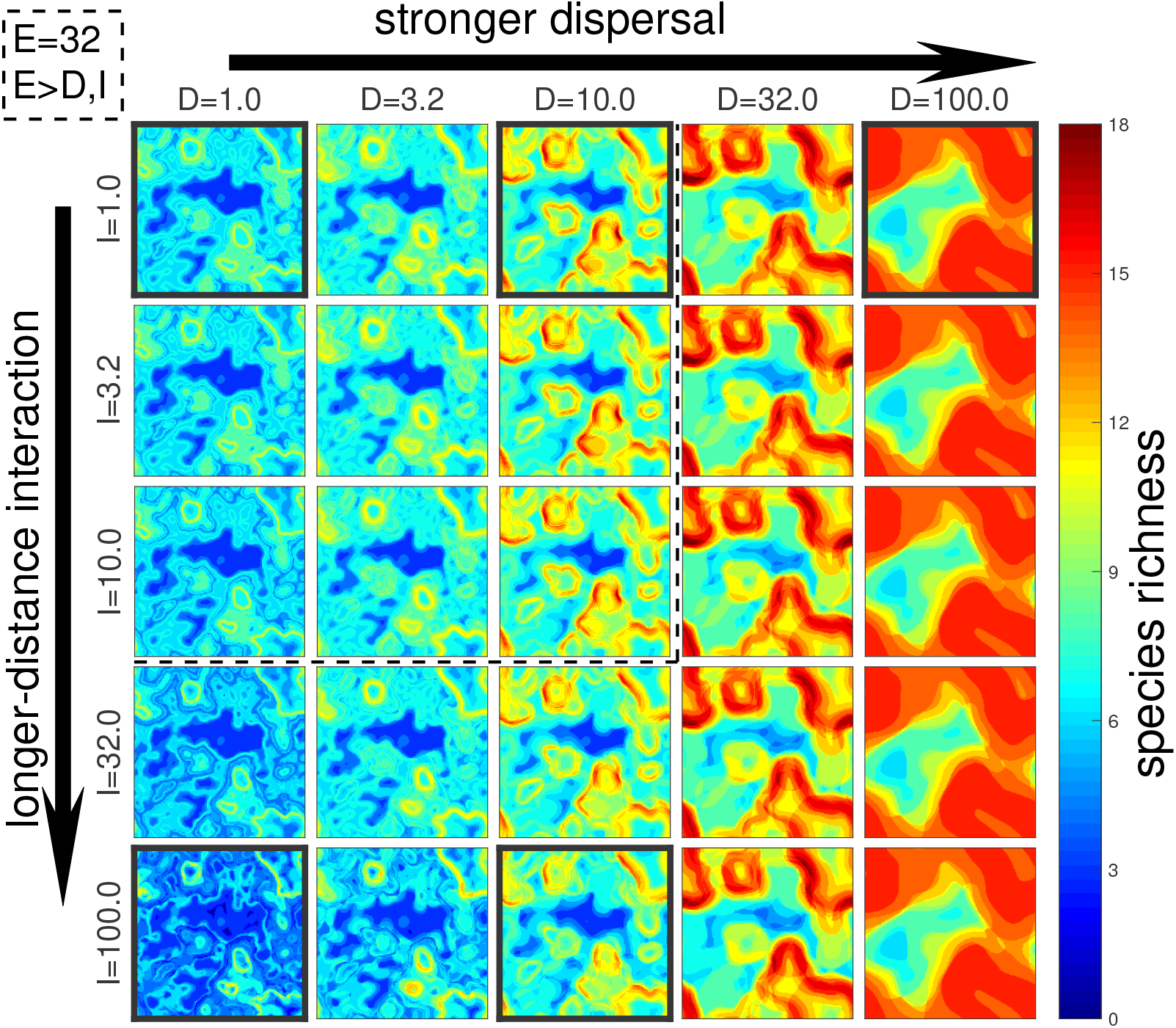
Species richness plots, corresponding to Fig.2, for the 5×5 scenarios (E = 32).

**Figure S10:**
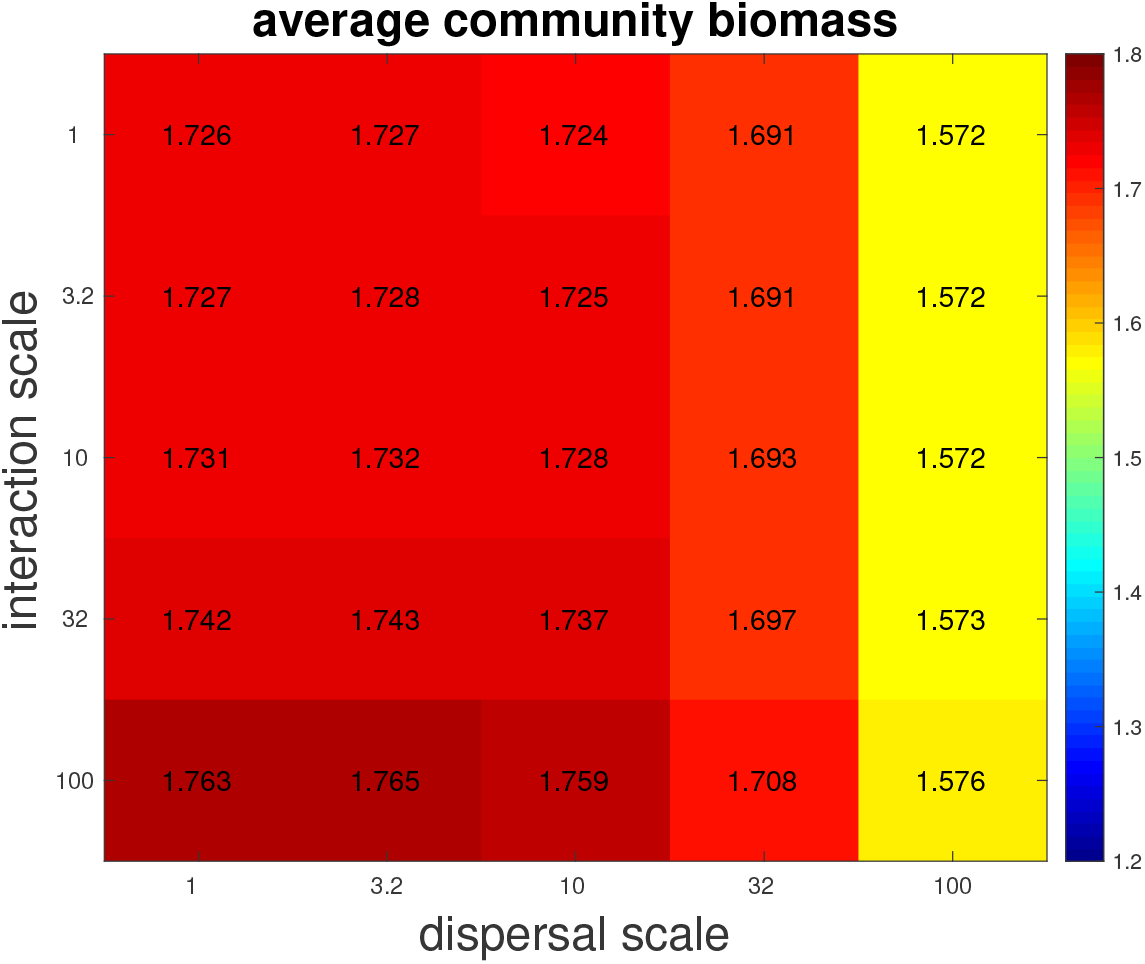
Total community biomass, averaged over domain, for the 5×5 scenarios (E = 32).

**Figure S11:**
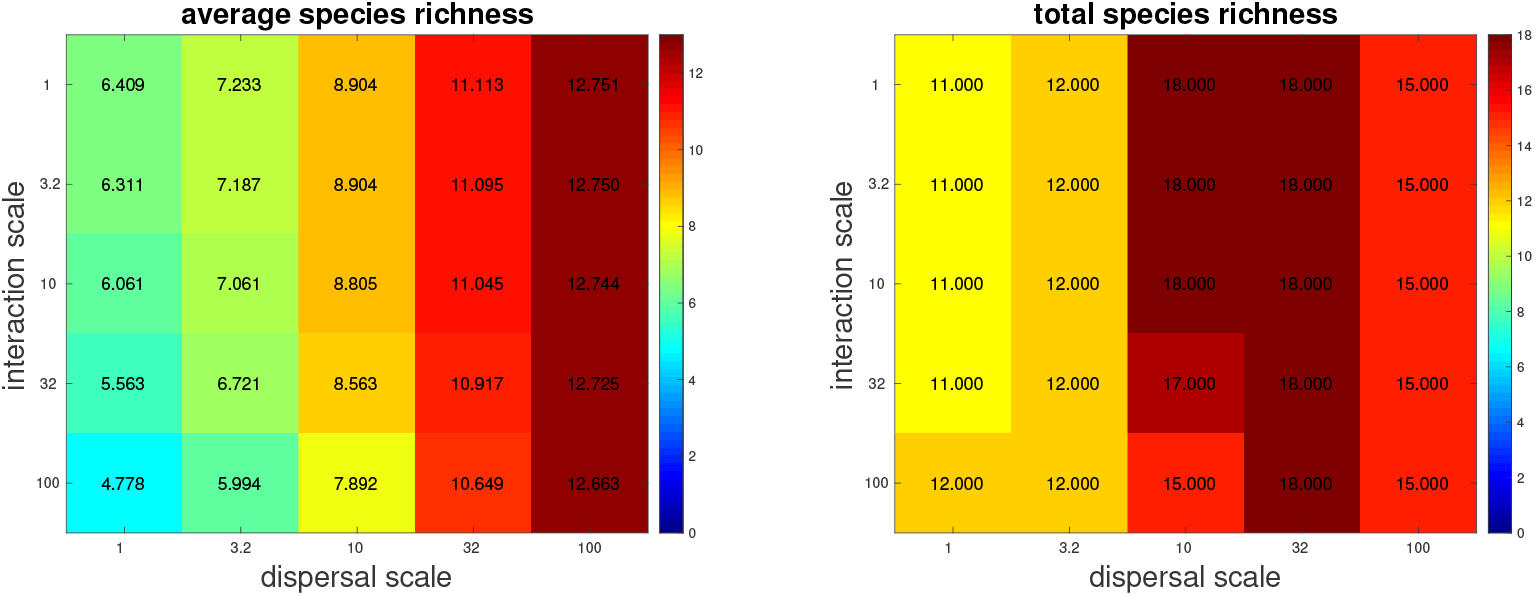
Diversity plots. Average local diversity of community (left) and total community diversity, (right) for the 5×5 scenarios (E = 32).

**Figure S12:**
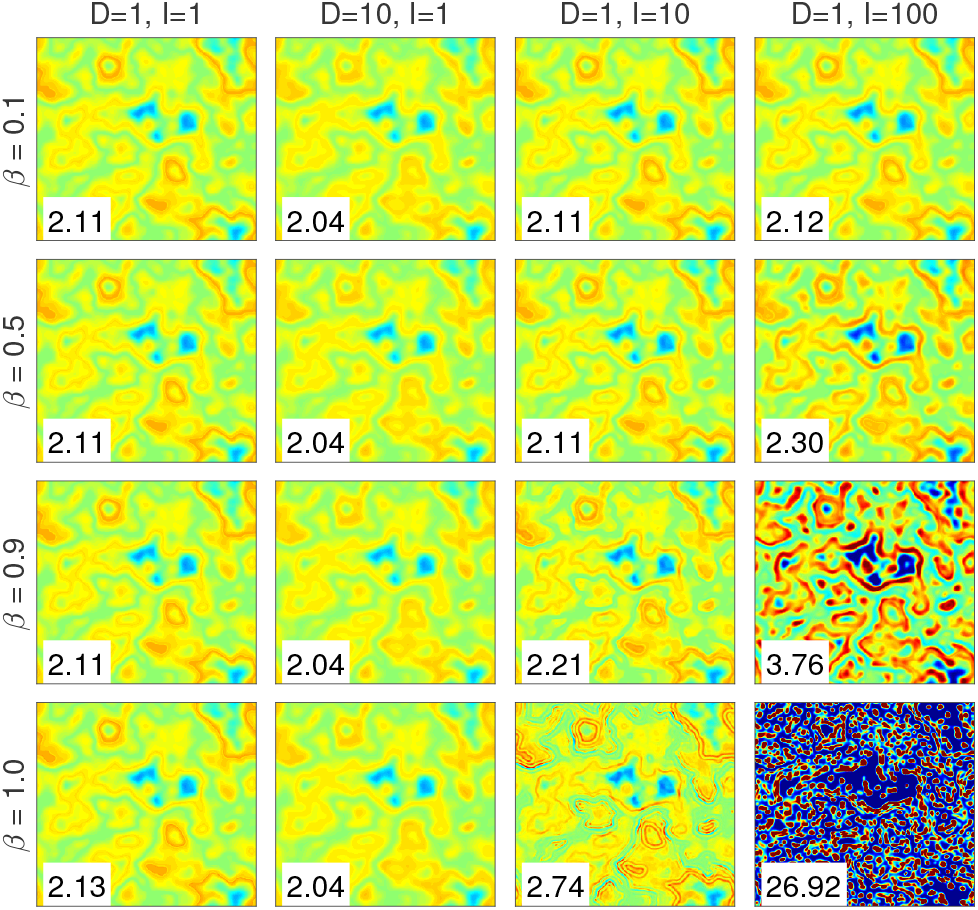
Effect of changing the value of the parameter. β, which determines the fraction of regional interactions. Each panel shows the spatial distribution of total biomass, with columns showing results for different values of I and D, while lower rows showing increasing values of β. The number in each panel shows the highest biomass density seen in the panel (where each panel’s colors are scaled to that value to better show the spatial structure). For low values of β (top two rows) scale of interactions I has minimal effect (clearly seen by right column looking the similar to other columns). For values of β (bottom two rows) the effect of I becomes strong and clearly visible. However, for very high values of β (bottom row) the effect also includes very high densities of biomass, which is not very realistic. We therefore choose a high value of β but not so high as to lead to very high densities (leading us to the middle ground of β = 0.9.

**Figure S13:**
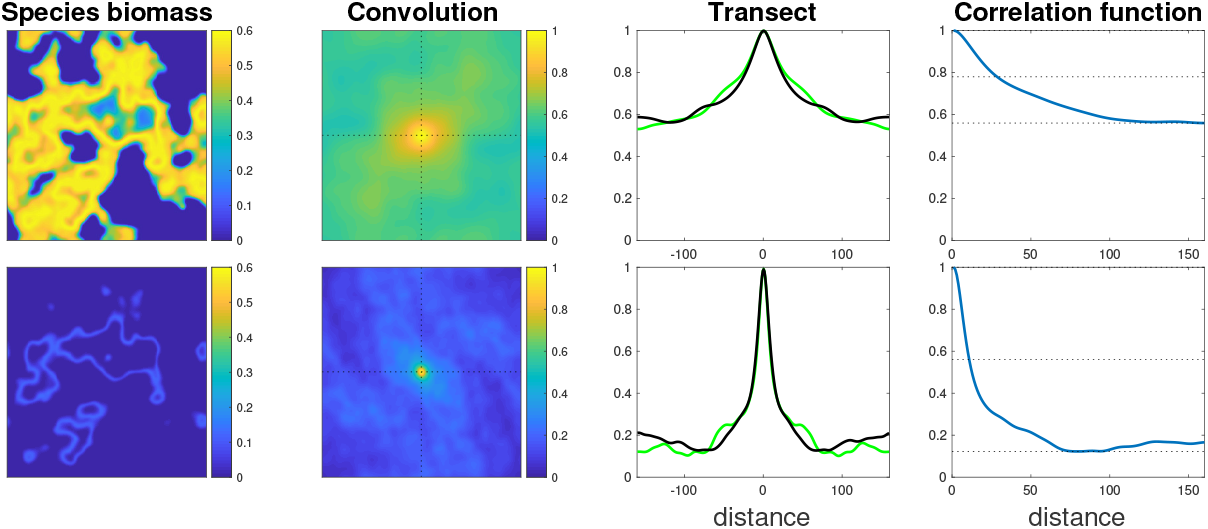
Demonstration of calculation of correlation function. The steps of calculating the correlation function are shown in the different columns, with the top (bottom) row corresponding to two different species in the same landscape used in Fig. 2. From left to right, the four columns correspond to: 1) The spatial distribution of biomass of a single species N_i_. 2) Correlation map, which is the result of a convolution of this spatial distribution with itself. 3) Transects of the correlation map (horizontal and vertical, shown in green and black), also marked in previous column by dotted lines. 4) Averaging of transects resulting in the correlation function. Horizontal dotted lines show the highest and lowest values of the correlation function, along with the average of the two which is used as a threshold to determine the scale of correlation.

